# Structural basis for transcript elongation control by NusG/RfaH universal regulators

**DOI:** 10.1101/324400

**Authors:** Jin Young Kang, Rachel Anne Mooney, Yuri Nedialkov, Jason Saba, Tatiana V. Mishanina, Irina Artsimovitch, Robert Landick, Seth A. Darst

## Abstract

NusG/RfaH/Spt5 transcription elongation factors are the only transcription regulators conserved across all life. In bacteria, NusG regulates RNA polymerase (RNAP) elongation complexes (ECs) across most genes, enhancing elongation by suppressing RNAP backtracking and also coordinating ρ-dependent termination and translation. RfaH is a specialized NusG paralog that engages the EC at *ops* sites and subsequently excludes NusG and suppresses both backtrack and hairpin-stabilized pausing. We used single-particle cryo-EM to determine structures of ECs at *ops* with NusG or RfaH. Both factors chaperone base pairing of the EC upstream duplex DNA to suppress backtracking. RfaH loads onto the EC by specific recognition of an *ops* hairpin in the single-stranded nontemplate DNA. Binding of both NusG and RfaH is incompatible with the swiveled RNAP conformation necessary for hairpin-stabilized pausing, but only RfaH fully counteracts swiveling to suppress pausing. The universal conservation of NusG/RfaH/Spt5 suggests that the molecular mechanisms uncovered here are widespread.

## INTRODUCTION

Multi-subunit RNA polymerases (RNAPs) interact with a wide array of accessory proteins that modulate every step of RNA synthesis. Among them, NusG/Spt5 is the only regulator that is conserved in all domains of life (Werner, 2012). NusG/Spt5 co-localizes with elongating RNAP across most genes (Mayer et al., 2010; Mooney et al., 2009a) and enhances transcript elongation by reducing RNAP pausing (Guo et al., 2000; Herbert et al., 2010; Hirtreiter et al., 2010). NusG/Spt5 may also attenuate transcription by cooperating with the negative regulator NELF to stimulate promoter-proximal pausing (Wu et al., 2003; Yamaguchi et al., 1999), by stimulating ρ-dependent termination (Peters et al., 2012), or by increasing pausing at certain sequences (Czyz et al., 2014; Sevostyanova and Artsimovitch, 2010; Yakhnin and Babitzke, 2014). NusG/Spt5 connects transcription to diverse cellular processes by bridging between RNAP and capping enzyme (Mandal et al., 2004), histone modifiers (Wier et al., 2013), somatic hypermutators (Pavri et al., 2010) and, in bacteria, ribosomes (Burmann et al., 2010). Specialized NusG/Spt5 paralogs have been identified in bacteria (Goodson et al., 2017), ciliates (Gruchota et al., 2017), and plants (Bies-Etheve et al., 2009). Specialized activities likely arose evolutionarily during organismal diversification; regulation of elongation via RNAP contacts the likely ancient and shared NusG/Spt5 activity.

NusG is a two-domain monomer in bacteria. Spt5 forms a heterodimer with Spt4 in archaea and eukaryotes (Hartzog and Fu, 2013). The NusG N-terminal domain (NGN), present in all NusG/Spt5-family proteins, contacts RNAP and is followed by KOW (Kyrpides et al., 1996) domains (one in bacteria and archaea; five in eukaryotes) and a C-terminal phosphorylated repeat region followed by two more KOW domains in eukaryotes. The NGN contacts the β’ clamp helices (CH) and the β gate loop (GL) on opposite sides of the active site cleft in RNAPs from all domains of life (Bernecky et al., 2017; Ehara et al., 2017; Klein et al., 2011; Sevostyanova et al., 2011; 2008), although an alternative location has been proposed based on a NusG-RNAP co-crystal structure lacking nucleic acid (Liu and Steitz, 2017). By bridging the cleft, the NGN has been proposed to function as a processivity clamp that ensures uninterrupted RNA synthesis. The NGN alone modulates RNAP pausing (Belogurov et al., 2007; Hirtreiter et al., 2010; Mooney et al., 2009b). The KOWs serve as contact sites for interacting proteins, but may also contact RNA or DNA (Bernecky et al., 2017; Ehara et al., 2017; Guo et al., 2015; Meyer et al., 2015) to aid elongation or stabilize binding to the EC (Crickard et al., 2016).

Life’s only universal transcription factor plays a central role in pausing, underscoring the importance of pausing in gene regulation. Pausing is proposed to aid timely recruitment of transcription regulators, guide nascent RNA folding, oppose chromatin silencing and genome instability, permit termination, and match the rate of transcription to those of other coupled processes, such as translation and splicing (Mayer et al., 2017). However, excessive pausing may lead to arrest, particularly in protein-coated DNA such as eukaryotic chromatin, which likely renders RNAP more susceptible to backtracking and imposes greater demands on processivity. Indeed, yeast Spt5 assists RNAP progression through the nucleosome (Crickard et al., 2017). Although recent results have suggested biophysical mechanisms for pausing via uncoupling of RNA and DNA translocation and for pause stabilization via rotation of an RNAP “swivel module” (Kang et al., 2018), the mechanism by which NusG/Spt5 suppresses pausing is unclear.

*E. coli* (*Eco*) RfaH is a NusG/Spt5 paralog that does not stimulate ρ, but exhibits strong anti-backtracking activity (like NusG/Spt5), can recruit ribosomes to nascent RNAs lacking a Shine-Dalgarno sequence via its KOW, and can counteract the pause-stabilizing effects of nascent RNA hairpins (pause hairpins; PHs) (Kolb et al., 2014). RfaH is recruited early in transcription units containing the *ops* sequence, which is exposed in the non-template strand DNA (nt-DNA) of paused elongation complexes (PECs) (Artsimovitch and Landick, 2002).

To investigate how interactions of NusG/Spt5/RfaH suppress backtrack and PH-stabilized pausing, and to visualize sequence-specific interaction of RfaH with the nt-DNA of ECs, we determined single-particle cryo-electron microscopy (cryo-EM) structures of *Eco* NusG or RfaH bound to an *ops*-containing EC (*ops*EC). We used the structures to design and interpret biochemical experiments that probe the interactions of RfaH and NusG with ECs and the mechanism by which they modulate pausing. Together, our results suggest a molecular model for the effects of NusG/Spt5-family proteins on transcription elongation.

## RESULTS

### Cryo-EM structures of a NusG-*ops*EC and an RfaH-*ops*EC

For cryo-EM structure determination of the NusG-*ops*EC and RfaH-*ops*EC, we designed an RNA-DNA scaffold based on the A20 (20mer RNA transcript with A at the 3’-end) scaffold used previously for cryo-EM structure determination of an *Eco* EC (Kang et al., 2017) except containing the *ops* sequence in the nt-DNA (Figure 1A). NusG suppresses pausing at and downstream from *ops*, whereas RfaH induces a strong pause 1-2 nucleotides after the *ops* pause (A20 on the *ops*EC scaffold; Figure 1A) (Artsimovitch and Landick, 2000; 2002). To ascertain that the *ops*EC obtained by direct reconstitution on the *ops*-scaffold supports NusG and RfaH function, we monitored RNA extension of radiolabeled A20 RNA in the presence of NusG or RfaH. In agreement with data obtained on standard templates, NusG reduced RNAP pausing downstream of *ops* whereas RfaH inhibited escape due to specific *ops*/RfaH interactions (Figure S1). We conclude that NusG or RfaH bind the directly reconstituted *ops*EC and modulate its function as expected.

**FIGURE 1.**
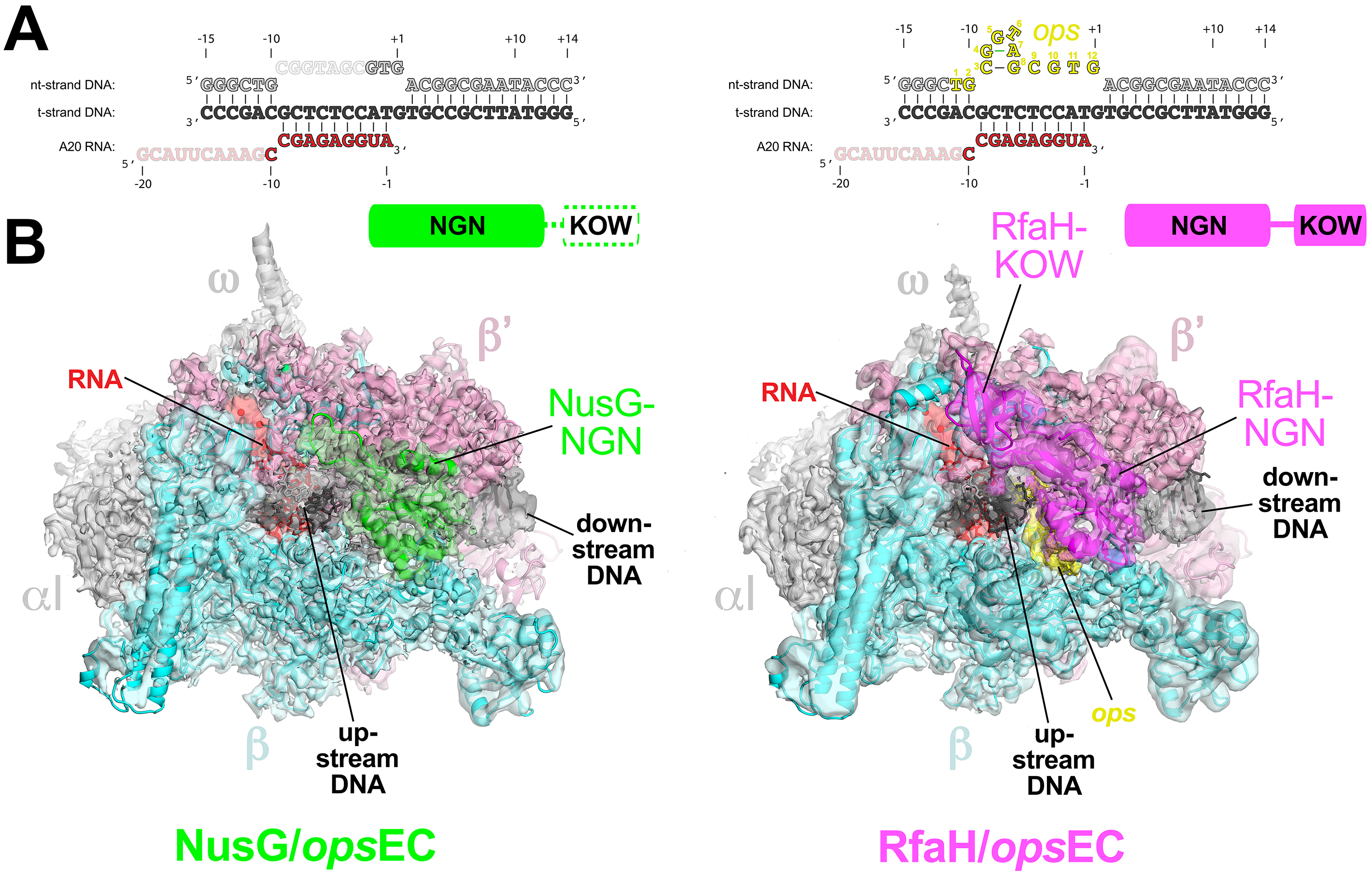
Structures of the NusG-*ops*EC and RfaH-*ops*EC. A. Nucleic acid scaffold sequence used for cryo-EM experiments. The same sequences are used in the NusG-*ops*EC (left) and RfaH-*ops*EC (right) structures. Disordered segments in each structure are faded. The nt-DNA *ops* sequence (nt-DNA −11 to +1, colored yellow for the RfaH-*ops*EC and numbered according to the *ops* position) forms a short hairpin that interacts specifically with RfaH but was mostly disordered in the NusG-*ops*EC structure (left). B. The cryo-EM density maps for the NusG-*ops*EC (left, 3.7 Å nominal resolution, low-pass filtered to the local resolution) and RfaH-*ops*EC (right, 3.5 Å nominal resolution, but shown is the 3.7 Å nominal resolution map with full-length RfaH, low-pass filtered to the local resolution) are rendered as transparent surfaces and colored as labeled. Superimposed are the final refined models; proteins are shown as backbone ribbons, the nucleic acids are shown as sticks. The schematics above indicate the domain organization of NusG (green) and RfaH (magenta). The NusG-KOW domain is disordered and is shown in white with a dashed green outline.

We used cryo-EM to determine the *Eco* NusG-*ops*EC and RfaH-*ops*EC structures (Figure 1B). The NusG-*ops*EC structure was determined to a nominal resolution of 3.7 Å (Figure 1B, S2, S3). Local resolution calculations indicate that the central core of the structure was determined to 3.0 - 3.5 Å resolution (Figure S3F). Although full-length NusG was used, cryo-EM density for only the NusG-NGN was observed; most of the NGN-KOW linker and the NusG-KOW were disordered (Figure 1B). Density corresponding to the NusG-KOW could not be recovered by focused classification approaches (Scheres, 2012)(see below).

The RfaH-*ops*EC structure was determined to a nominal resolution of 3.5 Å (Figures S4, S5). Local resolution calculations indicate that the central core of the structure was determined to 2.9 - 3.5 Å resolution (Figure S5G). Although full-length RfaH was used, cryo-EM density for only the RfaH-NGN was observed in the 3.5 Å resolution map. A particle classification focused on the flap-tip, RNA exit channel, and upstream duplex DNA gave rise to a second RfaH-*ops*EC reconstruction from a subpopulation of the particles (3.7 Å nominal resolution) that revealed cryo-EM density corresponding to the RfaH-KOW (Figures 1B, S4).

Initial examination of the structures revealed several key observations. First, both NusG and RfaH-NGN bound to the same location on the upstream face of the EC cleft, covering the single-stranded nt-DNA and upstream fork junction of the transcription bubble (Figure 1B). The NusG and RfaH-bound *ops*EC structures were similar to a previously reported *Eco* EC structure, also determined by cryo-EM (Kang et al., 2017), with rmsds of 0.67 Å (2,683 Cα′s aligned) and 0.648 Å (2,745 Cα’s aligned), respectively. The binding location and orientation of the NGN domains is consistent with biochemical analyses of NusG and RfaH interactions with the EC (Belogurov et al., 2007; 2010; Mooney et al., 2009b) as well as structural analyses of archaeal and metazoan Spt4/5 complexes (Bernecky et al., 2017; Ehara et al., 2017; Martinez-Rucobo et al., 2011) (Figures S6A, B). Our structures are not consistent with a crystal structure of an *Eco* NusG/core RNAP complex (Liu and Steitz, 2017) but proper binding of the NusG-NGN is precluded by neighboring symmetry-related RNAP molecules in this crystal lattice, explaining the discrepancy (Figure S6C).

Second, binding of the NusG- and RfaH-NGNs remodel the channels for the upstream duplex DNA, the upstream fork-junction of the transcription bubble, and the upstream segment of the single-stranded nt-DNA, stabilizing the upstream duplex DNA and possibly explaining the suppression of backtrack pausing (Figure 2). In the RfaH-*ops*EC, the *ops* sequence in the single-stranded nt-DNA forms a short hairpin-like structure that interacts sequence-specifically with RfaH (Figure 3A). Space exists in the NusG-*ops*EC to accommodate the nt-DNA but NusG lacks the capacity to interact sequence-specifically with the DNA and much of the *ops* sequence in this context is disordered (Figure 1A).

**FIGURE 2.**
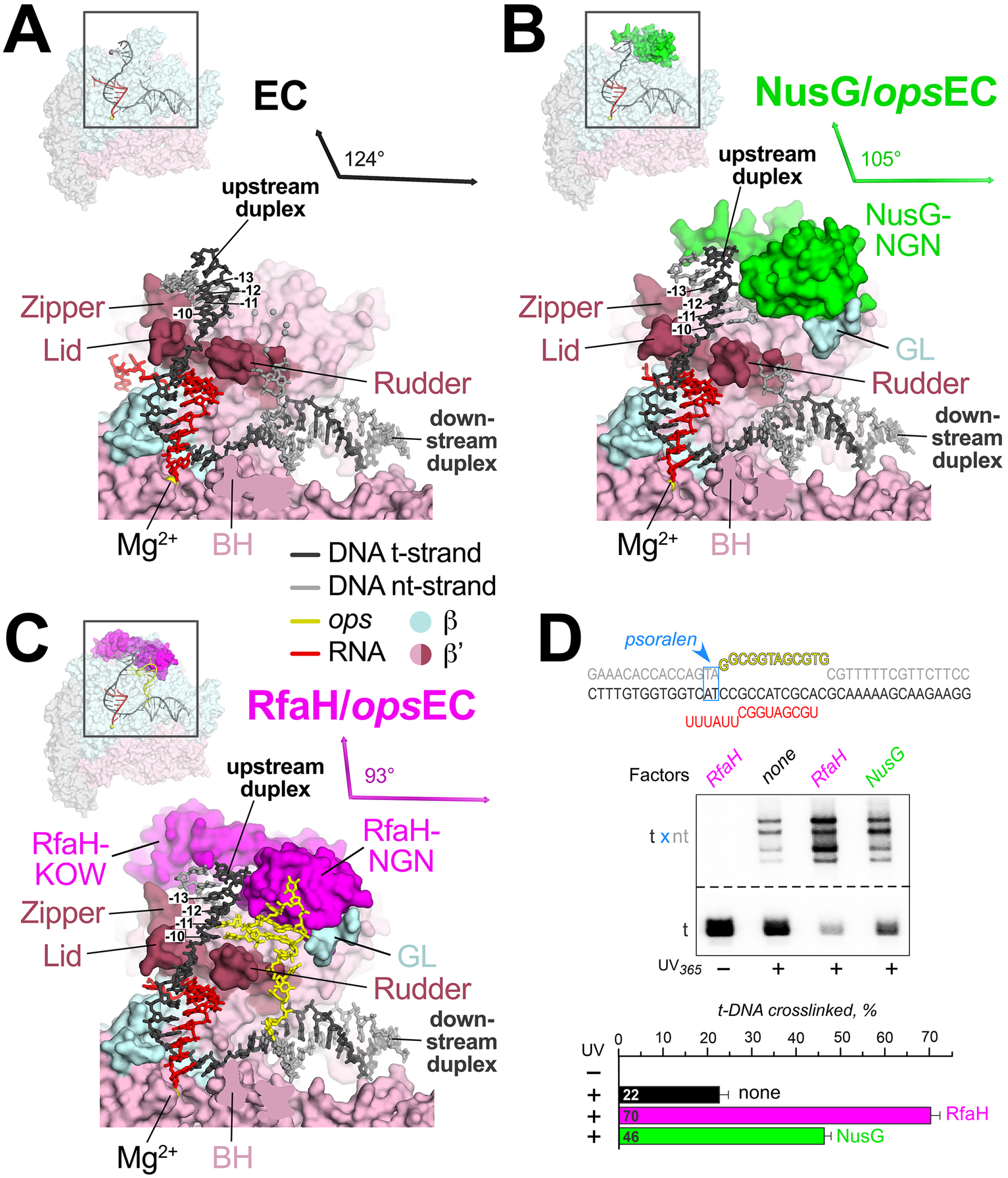
NusG and RfaH remodel and stabilize the upstream duplex DNA. A. (top left) Overall view of the EC structure (Kang et al., 2017). The RNAP is shown as a transparent molecular surface, revealing the nucleic acid scaffold inside (shown in cartoon format and colored according to the legend). The boxed region is magnified below. (bottom) Magnified view of boxed region from overall view above. Most of the β subunit (light cyan) has been removed to reveal the inside of the RNAP active site cleft. The β’ subunit is light pink but with the zipper, lid, and rudder highlighted in brown. The Bridge-Helix (BH) is also labeled. The nucleic acids are shown as sticks (with the first four base pairs of the upstream duplex, −10 through −13, labeled). The RNAP active site Mg^2+^-ion is shown as a yellow sphere. The thin black arrows are drawn parallel to the downstream duplex DNA axis (nearly horizontal) and the upstream duplex (Lu and Olson, 2008), which subtends an angle of 124°. B. As in (A) but showing the NusG-*ops*EC structure. NusG is colored green. C. As in (A) but showing the RfaH-*ops*EC structure. RfaH is colored magenta. D. Probing the upstream fork junction by psoralen crosslinking. The *ops*ECs were assembled on the scaffold shown on top, with the TA intercalation motif (blue) positioned immediately upstream from the *ops* element; the t-DNA was labeled with [γ^32^P]-ATP. Following incubation with RfaH or NusG, the ECs were illuminated with 365 nm UV light. The crosslinked products were analyzed on 12 % gels and the fraction of t-DNA crosslinked to the nt-DNA was quantified (bottom). Error bars indicate the s.d. of triplicate measurements.

**FIGURE 3.**
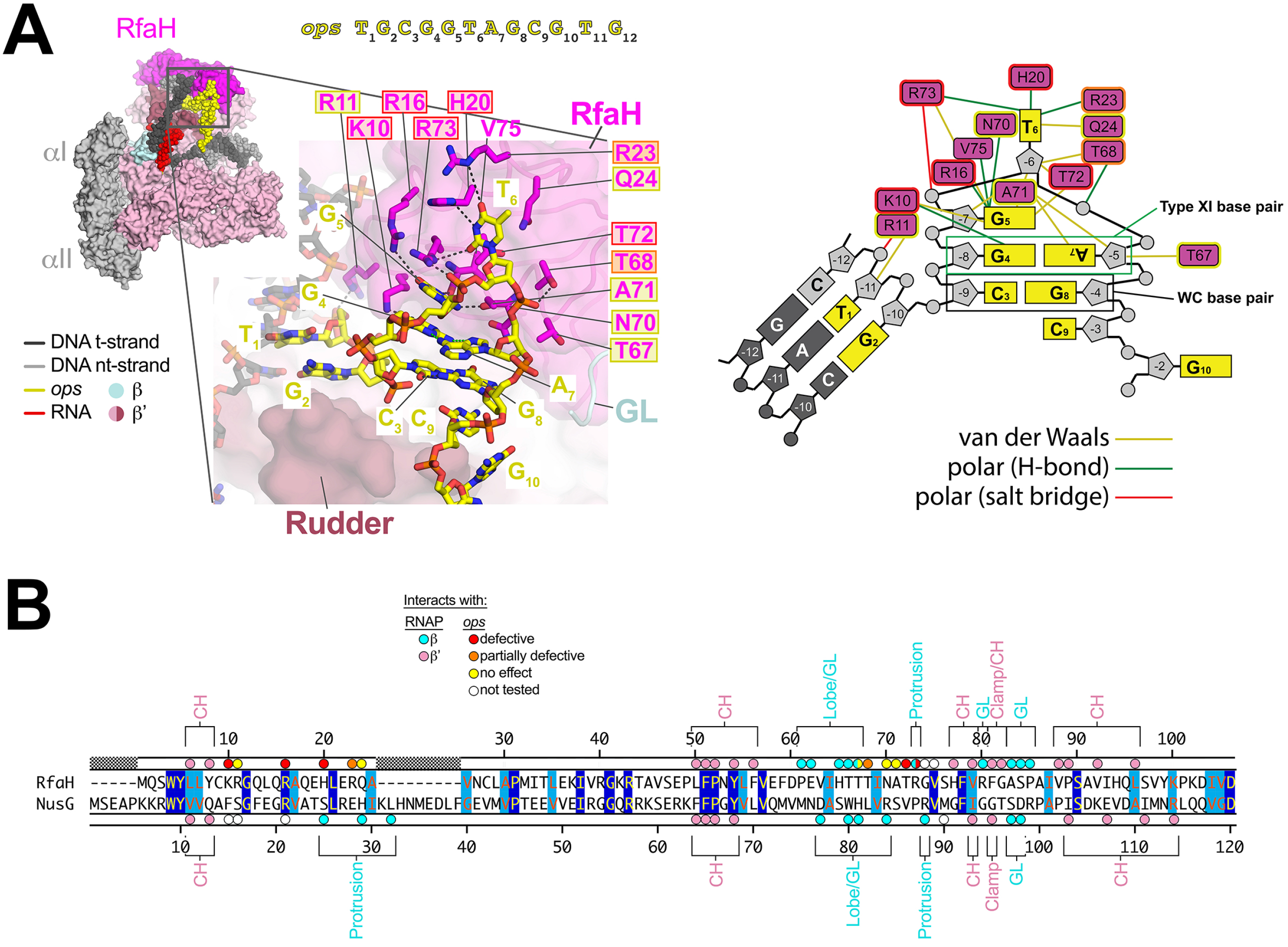
RfaH/*ops* interactions. A. (left panel, top) View of the RfaH-*ops*EC (similar to the view of Figure 2C); the nt-DNA sequence shown on top is numbered according to *ops* sequence position. Proteins are shown as molecular surfaces. The nucleic acids are shown in CPK format and colored according to the legend. The boxed region is magnified with RfaH rendered transparent, revealing the α-carbon backbone (in cartoon format) and amino acid side chains that interact with *ops*. Polar RfaH/*ops* interactions, H-bonds (≤ 3.5 Å) or salt bridges (≤ 4.5 Å) are denoted by gray dashed lines. The *ops* sequence (yellow) is labeled and numbered. The RfaH residues are labeled. The shaded boxes denote the effect of substitutions on RfaH recruitment to *ops* (red shaded box, defective; orange shaded box, partially defective; yellow shaded box, no effect; no box, not tested; (Belogurov et al., 2010). (right panel) Schematic representation of RfaH/*ops* interactions. The DNA is color-coded as in Figure 3A. The magenta rectangles denote RfaH residues contacting the DNA. Colored lines denote interactions: yellow, van der Waals (≤ 4.5 Å); green, H-bond (≤ 3.5 Å); red, salt bridge (≤ 4.5 Å). B. Structure-based sequence alignment of the *Eco* RfaH-NGN and NusG-NGN, with residues numbered above and below, respectively. Identical residues are shaded dark blue, homologous residues light blue. The colored dots on top (for RfaH) and bottom (for NusG) denote RNAP and *ops* contacts (color-coded as shown in the legend). The RNAP structural elements that the RfaH (top) and NusG (bottom) residues interact with are denoted.

Third, the NusG-and RfaH-NGNs make contacts with the RNAP that bridge across the upstream face of the active site cleft (Figures 4A, B), stabilizing the overall active conformation of the EC and disfavoring the swiveled conformation associated with PH-stabilized pausing (Figure 4C) (Kang et al., 2018). The stabilization of the active EC RNAP conformation by RfaH is much stronger than NusG (Figure 5A-F), and RfaH is much more effective at inhibiting hairpin-stabilized pauses than NusG (Figure 5C) (Artsimovitch and Landick, 2000; 2002; Belogurov et al., 2009). We will describe these structural features in the context of the roles of NusG and RfaH in transcription elongation, and present biochemical evidence for their roles.

**FIGURE 4.**
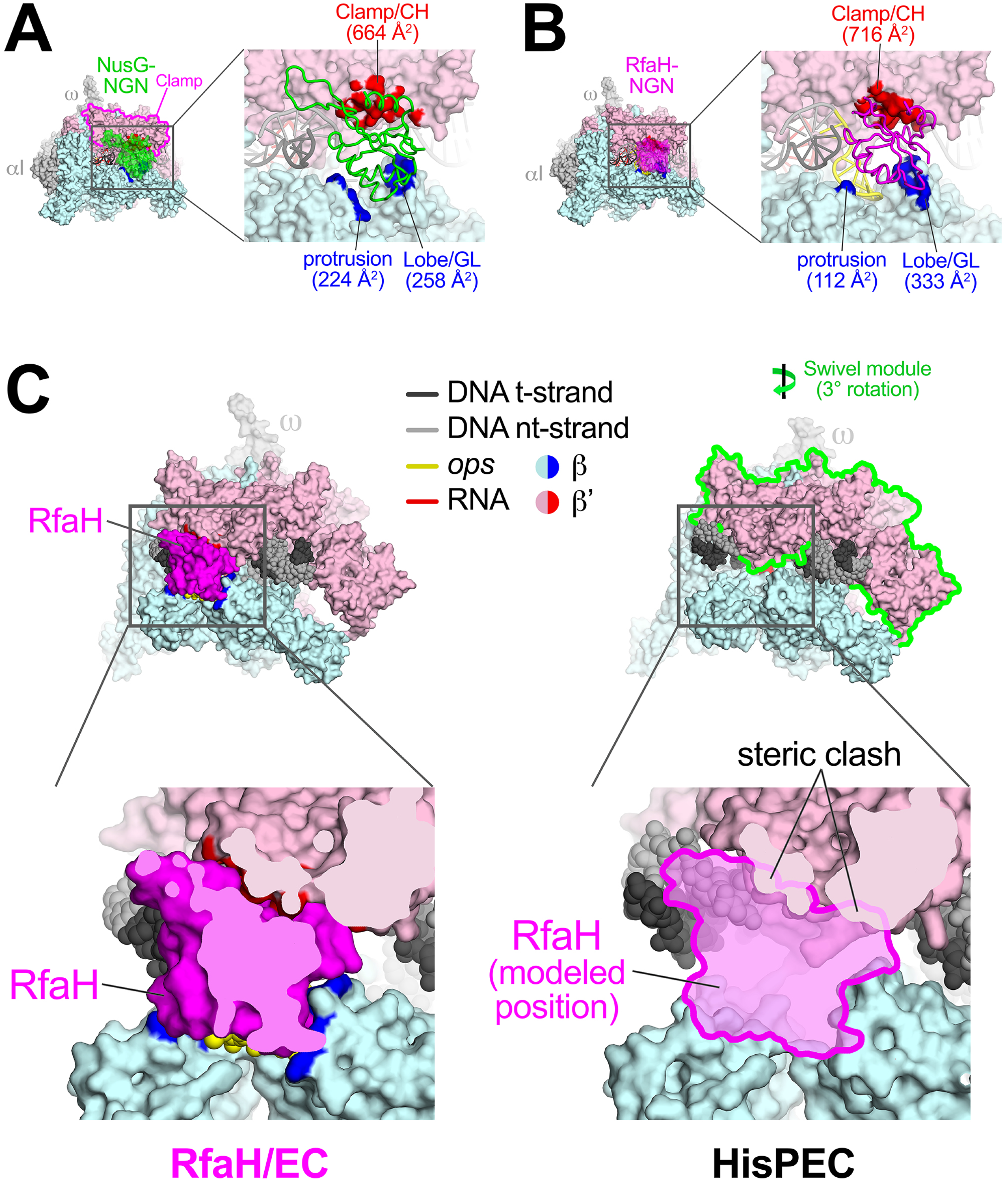
NusG-NGN and RfaH-NGN interactions with RNAP bridge the active site cleft and are incompatible with PH-induced RNAP swiveling. A. (left) Overall view of the NusG-*ops*EC structure. Proteins are shown as molecular surfaces (color-coded as shown in the key or as labeled). The nucleic acids are shown in cartoon format and color-coded as shown in the key. (right) Magnified view of the boxed region on the left. The NusG-NGN is shown as an a-carbon backbone worm. Surfaces of RNAP that contact NusG (≤ 4.5 Å) are colored red (β’ subunit) or blue (β subunit) and labeled along with the buried surface area of the protein/protein interaction. B. Same as (A) but for the RfaH-*ops*EC structure. C. (left, top) Overall view of the RfaH-*ops*EC structure. The boxed region is magnified below. (left, bottom) Magnified view of the boxed region from above, sliced at the level of RfaH to reveal the close fit between RfaH and elements of the RNAP β (light cyan) and β’ (light pink) subunits. (right, top) Overall view of the *his* PH-stabilized PEC [Kang et al., 2018]. The formation of the PH in the RNAP RNA exit channel induces an ~3° rotation (as shown) of the ‘swivel module’ (outlined in green; Kang et al., 2018). The boxed region is magnified below. (right, bottom) Magnified view of the boxed region from above, but also showing the modeled position of RfaH. The swiveled conformation of the *his*PEC is not compatible with RfaH binding due to steric clashes (denoted).

**FIGURE 5.**
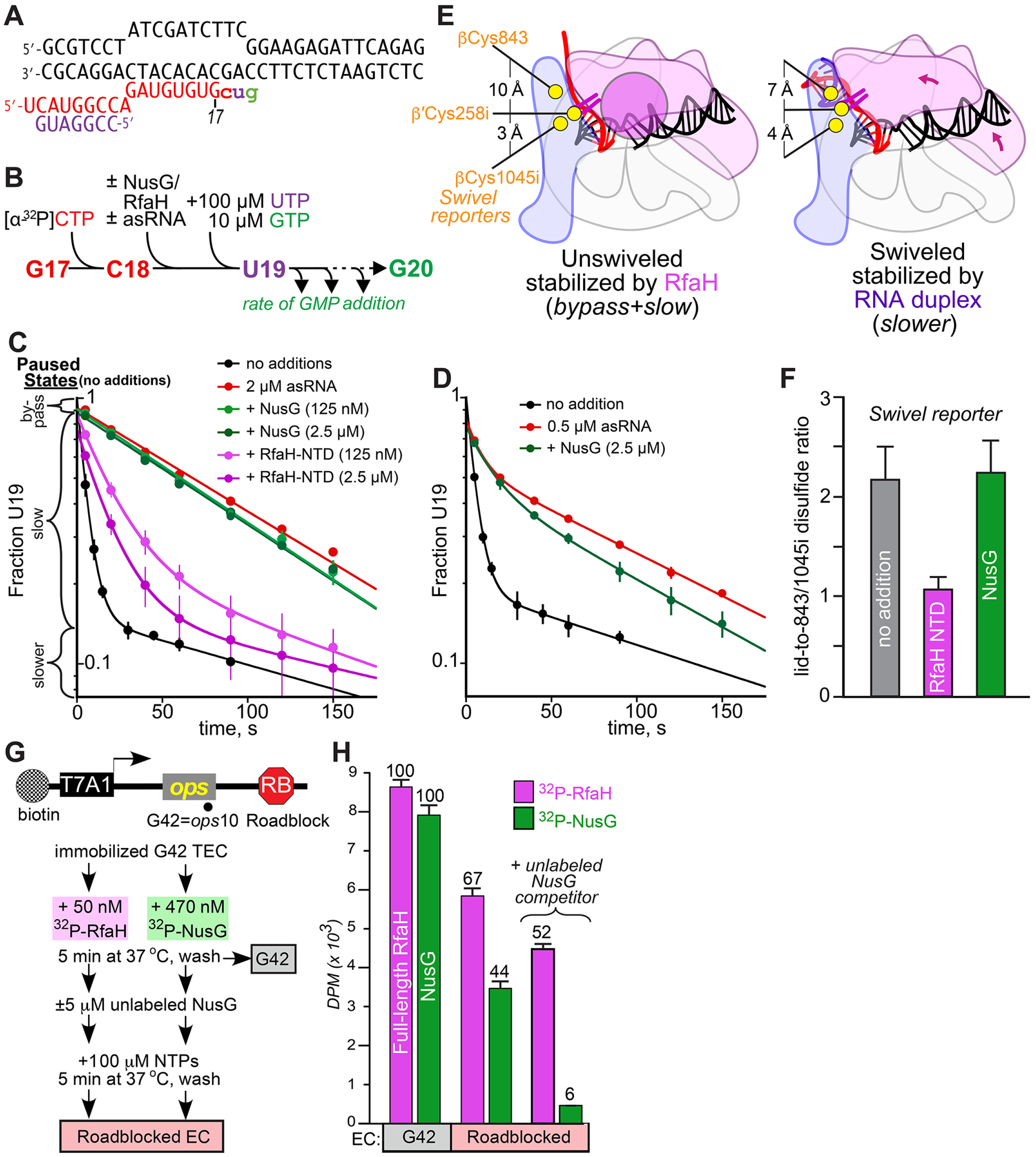
RfaH inhibits PH stimulation of pausing and RNAP swiveling more effectively and binds ECs more tightly than NusG. A. Scaffold used to test asRNA stimulation of pausing. The sequence is identical to that used to determine the cryo-EM structure of the *his*PEC (Kang et al., 2018), except that the PH is replaced with a target for asRNA binding (Kolb et al., 2014). B. Experimental scheme to measure PH stimulation of pausing using asRNA. C. Effect of RfaH-NGN and NusG on stimulation of pausing by 2 μM asRNA. The fraction of U19 present as a function of time after UTP and GTP addition was triphasic. A small fraction of ECs failed to enter the paused state (bypass fraction). In the absence of asRNA, most U19 entered the elemental paused state and exhibited slow addition of G20. In the absence of asRNA, a small fraction of U19 ECs added G20 more slowly and may reflect formation of the swiveled EC. RfaH-NGN, but not NusG, inhibited formation of the slow fraction. Data shown are means and s.d. from at least three replicates. D. Effect of NusG on stimulation of pausing by 0.5 μM asRNA. NusG decreased the slower, apparently swiveled fraction of U19 only modestly, from 0.57 ± 0.02 to 0.49 ± 0.02. Data shown are means and s.d. from at least three replicates. E. Graphical depiction of RNAP swiveling and location of CTR Cys residues. PH-induced swiveling changes distance from β′C258i to its potential disulfide partners, βC1045i or β’C843 from 3 Å and 10 Å, respectively, to 4 Å and 7 Å (Kang et al., 2018). F. RfaH-NGN but not NusG decreased the ratio of CTR reporter disulfides on the *his*PEC scaffold (containing PH; Kang et al., 2018), indicating that RfaH-NGN but not NusG shifts the equilibrium from the swiveled toward the unswiveled RNAP conformation. Data shown are means and s.d. from at least three replicates. G. Measuring RfaH and NusG retention on the ECs. A linear DNA template with a T7A1 promoter, the *ops* element, and an EcoRI site 128 nt downstream from the *opsP* site was immobilized on streptavidin beads via a biotin on the nt-DNA (top). A cleavage-deficient EcoRI^Q111^ protein (RB) was bound to roadblock the transcribing RNAP. Halted *ops*10 (G42) ECs were formed by step-wise transcription with NTP subsets and incubated with radiolabeled RfaH or NusG. After washing away the unbound RfaH/NusG, transcription was resumed by addition of all NTPs with or without an excess of unlabeled NusG, which is expected to bind to RNAP upon dissociation of the pre-bound factor. G42 and roadblocked ECs were washed to remove unbound RfaH/NusG and analyzed by scintillation counting. H. RfaH (magenta bars) and NusG (green bars) binding to the immobilized ECs. The residual factor binding to the roadblocked ECs is expressed relative to that observed with the G42 EC, which is defined as 100%. Error bars indicate the s.d. of triplicate measurements.

### NusG and RfaH remodel the EC nucleic acids, chaperoning upstream duplex DNA reannealing and explaining the suppression of backtrack pausing

After separating from the RNA-DNA hybrid, the template strand DNA (t-DNA) in the EC is directed out of the RNAP active site cleft through a channel between the β’lid and the β’rudder where it immediately anneals with the nt-DNA (-10 position, Figures 1A, 2A) (Kang et al., 2017). The upstream DNA duplex is relatively unconstrained and mobile, making few interactions with the RNAP (Kang et al., 2017; Korzheva et al., 2000).

Other than RfaH-R11, which interacts with the nt-DNA phosphate backbone at the −11 position (*ops* T_1_; Figure 3A), stable interactions between NusG or RfaH and upstream duplex DNA are not observed and consequently the DNA segment is mobile. Nevertheless, both NusG and RfaH remodel the path of the upstream duplex DNA, decreasing the subtended angle with the downstream duplex DNA (Figures 2B, C).

Psoralen intercalates into double-stranded DNA at TA steps and forms a T-T interstrand crosslink when activated by UV light. Psoralen crosslinking efficiency serves as a probe of DNA structure since psoralen intercalates most efficiently in stable, B-form duplex DNA. Based on results of psoralen crosslinking and fluorescence quenching, (Turtola and Belogurov, 2016) proposed that the −10 base pair at the upstream fork-junction of the EC transcription bubble was distorted, as later observed in the EC structure (Kang et al., 2017). We used the same approach to probe the upstream fork-junction of the *ops*EC. NusG increased the efficiency of crosslinking at the upstream fork-junction more than two-fold over the EC (Figure 2D), as observed previously (Turtola and Belogurov, 2016). RfaH increased the crosslinking efficiency more than 3.5-fold over the EC. Taken together, these results suggest that NusG and RfaH chaperone and stabilize the formation of the −10 bp.

### RfaH recognizes *ops* as a nt-DNA hairpin

In the EC, the 10 nucleotides of the single-stranded nt-DNA (-9 to +1) in the transcription bubble span from the −10 to +2 nucleotides, which are base-paired in the upstream and downstream DNA duplexes, respectively (Figure 1A). Repositioning of the upstream duplex DNA by NusG/RfaH significantly reduces the distance separating the −10 and +2 nt-DNA phosphates from 41 Å in the EC to 33 Å in the NusG-*ops*EC and 30 Å in the RfaH-*ops*EC. In the RfaH-*ops*EC, a short hairpin with a two base pair stem forms in the single-stranded nt-DNA and this DNA structure is specifically recognized by RfaH (Figure 3A).

Using the *ops* sequence numbering (Figure 3A) rather than the scaffold numbering, T_1_ and G_2_ of *ops* are base-paired as part of the upstream DNA duplex (Figure 3A). C_3_ forms a Watson-Crick base pair with G_8_, while G_4_ and A_7_ participate in a Saenger type XI base pair (Saenger, 1984) to form the *ops* hairpin stem. G_5_ stacks on the upstream face of G_4_ while T_6_ is flipped out of the base stack. C_9_ stacks with the downstream face of G_8_. The rest of the *ops* sequence (G_10_T_11_G_12_) is single-stranded and does not interact with RfaH but interacts with RNAP as in the EC structure. Most notably T_11_ stacks with βW183 and G_12_ binds in a G-specific pocket of the β subunit (Kang et al., 2017; Zhang et al., 2012).

Other than RfaH-K10, which hydrogen-bonds (H-bonds) with G_4_(O_6_), the base pairs C_3_:G_8_ and G_4_:A_7_ do not make extensive base-specific interactions with RfaH - these bases are conserved in the *ops* sequence because of their role in forming the *ops* hairpin stem. The geometry of the *ops* hairpin stem sets up extensive base-specific interactions of RfaH with G_5_ and T_6_ (Figure 3A). The three potential H-bonding atoms of the Watson-Crick edge of G_5_ (N_2_, N_1_, O_6_) H-bond with the backbone carbonyls of RfaH-N70 and V75, and with the side chain of R16, respectively. The three potential H-bonding atoms of the T_6_ base (O_2_, N_3_, O_4_) participate in H-bonds with the side chains of R73, H20, and R23, respectively. The aliphatic chain of RfaH-Q24 makes van der Waals contact with the T_6_ exocyclic methyl. All of the RfaH side chains that make base-specific H-bonds (K10, R16, H20, R23, R73) are important for RfaH function. Single Ala substitutions of these residues interfere with RfaH recruitment at *ops* (Figure 3) (Belogurov et al., 2010).

In the structure-based alignment with NusG, RfaH-K10, H20, R23, and R73 correspond to NusG-F15, S25, E28, and P87, each unable to participate in the equivalent interactions with *ops* (Figure 3B). The disposition of the DNA in the NusG-*ops*EC seems compatible with *ops* hairpin formation (Figure 2B) but cryo-EM density for most of the *ops* sequence (C_3_ to C_9_) is completely absent and the DNA is presumed to be disordered.

### NusG and RfaH contacts bridge the RNAP active-site cleft

The overall RNAP structure has been likened to a crab claw, with one pincer comprising primarily the β’ subunit, and the other primarily the β subunit (Figure 4) (Zhang et al., 1999). Between the two pincers is a large cleft that contains the active site and accommodates the nucleic acids in the EC (Gnatt et al., 2001; Kang et al., 2017; Korzheva et al., 2000; Vassylyev et al., 2007). The clamp (Figure 4A), a mobile structural module that makes up much of the β’ pincer (Gnatt et al., 2001), undergoes swinging motions that open the channel to allow entry of nucleic acids during initiation, or that close the channel around the DNA and RNA-DNA hybrid to enable processive elongation (Chakraborty et al., 2012; Feklistov et al., 2017; Gnatt et al., 2001).

The NusG and RfaH-NGN bind the upstream face of the EC, bridging the active-site cleft by making significant contacts with the clamp of the β’ pincer and the protrusion and lobe of the β pincer (Figures 4A, B). NusG and RfaH interactions with RNAP are analogous in that the same regions of each factor interact with the same regions of RNAP (Figure 3B) with one exception - the first α-helix of NusG (residues 18-34) interacts with the protrusion while the same region of RfaH (residues 13-24) interacts with the *ops* hairpin, which inserts between RfaH and the protrusion (Figure 4B). Despite these additional NusG/RNAP interactions, overall the NusG/RNAP and RfaH/RNAP interface buries a similar total surface area (NusG, 1,150 Å^2^; RfaH, 1,160 Å^2^). After RNAP escape from *ops*, the specific RfaH-*ops* contacts are lost and RfaH may establish interactions with the protrusion, significantly increasing the RfaH/RNAP interaction interface and affinity relative to NusG. This is consistent with observations that RfaH outcompetes NusG for EC binding *in vitro* even when NusG-NGN is at a 10-fold excess over the RfaH (Belogurov et al., 2009), and excludes NusG from RNAP transcribing *ops*-operons in the cell (Belogurov et al., 2009) despite NusG being present in large excess (50 to 100-fold) over RfaH *in vivo* (Schmidt et al., 2016).

The primary interaction surface for both NusG and RfaH is with the clamp helices (CH; Figures 4A, B), consistent with previous analyses (Belogurov et al., 2007; 2010; Mooney et al., 2009b; Sevostyanova et al., 2008). Both factors interact with the GL. However, interactions of the RfaH HTT motif (residues 65-67; Figure 3B) with the GL are required for RfaH function (Belogurov et al., 2010; Sevostyanova et al., 2011), whereas deletion of the GL supports normal NusG activity (Nandymazumdar et al., 2016; Turtola and Belogurov, 2016).

### Both NusG and RfaH contacts are incompatible with RNA pause hairpin-induced EC swiveling

RfaH efficiently suppresses both backtrack-and PH-stabilized pausing (Artsimovitch and Landick, 2002; Kolb et al., 2014). Nascent PHs can increase pause lifetimes tenfold or more (Toulokhonov et al., 2001). Recent cryo-EM studies revealed that formation of the *his*PH in the RNAP RNA exit channel induced a previously unseen global conformational change in the RNAP termed ‘swiveling’ (Kang et al., 2018). In the swiveled RNAP, the clamp and other structural features of the β’ pincer (called the swivel module; Figure 4C) undergo a concerted rotation of about 3° about an axis roughly parallel with the BH (or perpendicular to the RNA-DNA hybrid). The PH-induced swiveling module is thought to increase pause lifetimes by allosterically inhibiting trigger-loop folding (Kang et al., 2018). Swiveling alters the relative positions of the β’ and β pincers which the bound RfaH bridges, and modeling reveals that RfaH binding is incompatible with the swiveled state (Figure 4C).

NusG binds to RNAP similarly (Figure 3B), bridging the β’ and β pincers of RNAP (Figure 4A), and modeling also indicates that NusG binding is incompatible with the swiveled conformation. The greater inhibition of PH action and of RNAP swiveling by RfaH is likely due to stronger binding of RfaH to ECs. To explore these differences, we used a scaffold resembling the *ops* cryo-EM scaffold but containing *his* pause sequences that form an RNA-duplex-stabilized pause upon addition of an antisense RNA oligonucleotide (asRNA; Figure 5A; Kang et al., 2018). RfaH, but not NusG, can outcompete binding of an 8mer asRNA to a similar scaffold (Kolb et al., 2014); thus, we used a 7mer asRNA to maximize the ability of NusG to compete (Figure 5A). When added to radiolabeled C18 complexes formed one nucleotide upstream of the pause, the 7mer asRNA stimulated pause dwell time at U19 ~20-fold (Figures 5B, C, E, slow fraction minus asRNA representing elemental PECs vs. slower fraction plus asRNA representing RNA-duplex-stabilized PECs; the bypass fraction represents ECs that failed to enter the pause; a small fraction of slower, swiveled PEC appeared to form even in the absence of asRNA). Whereas 125 nM RfaH could suppress pause stimulation by 2 μM asRNA almost as effectively as 2.5 μM RfaH, 2.5 μM NusG had little or no effect on asRNA action (Figure 5C). When asRNA was lowered to 0.5 μM, 2.5 μM NusG gave a minimally detectable effect (Figure 5D).

Consistent with an ability of RfaH but not NusG to suppress asRNA-induced swiveling, a cysteine-triplet reporter (CTR) that detects swiveling by a shift in disulfide bond formation by β′-lid Cys258i from βCys1045i to βCys843 (Kang et al., 2018) reported RfaH but not NusG suppression of the swiveled conformation on a *his*PEC scaffold (Figures 5E, F).

To ask if this reduced effect of NusG could be explained by weaker binding of NusG vs. RfaH to ECs, we performed a NusG-RfaH competition experiment (Figure 5G). Radiolabeled RfaH or NusG was bound to ECs halted at *ops* by step-wide waling of RNAP after initiation on a T7 A1 promoter template. Upon addition of all four NTPs, ECs moved along the template until encountering a roadblock generated by a noncleaving mutant *Eco*RI endonuclease bound downstream of *ops* (Pavco and Steege, 1990; Strobel et al., 2017). When the roadblocked ECs, in which *ops*-RfaH contacts were no longer possible, were washed with buffer, radiolabeled RfaH was retained to a greater extent than radiolabeled NusG (Figure 5H). Almost no radiolabled NusG was retained when unlabeled NusG competitor was present at 5 μM, whereas most radiolabeled RfaH remained bound in the presence of NusG competitor. These data establish that, once assocated with an EC at *ops*, RfaH remains bound even in the absence of *ops* contacts, whereas similarly bound NusG is readily lost, consistent with much weaker NusG-EC binding than RfaH-EC binding.

### RfaH KOW domain binds RNAP and remodels the upstream duplex DNA path and RNA exit channel like Spt5

The positioning of Spt5 KOW1-5 domains on the surface of RNAPII by cryo-EM (Bernecky et al., 2017; Ehara et al., 2017) suggests that the Spt5-KOWs may contact upstream DNA, RNA, and transcription regulators from fixed locations on the EC, rather than on freely rotating tethers. Thus, our finding that the RfaH-KOW binds RNAP at a location similar to that occupied by Spt5-KOW1 merits comparison (Figure 6). The side-chain contacts and exact positions of Spt5-KOW1 and RfaH-KOW are not conserved, but their locations relative to RNAP and upstream duplex DNA are similar. For both structures, the open face of the large KOW domain b sheet faces away from RNAP, but the RfaH-KOW is rotated ~45° and shifted about 10 Å toward the RNA exit channel relative to Spt5-KOW1. Both KOWs appear to guide the upstream duplex DNA.The RfaH-KOW shifts the duplex trajectory and narrows the angle with downstream DNA, whereas Spt5-KOW1 does not significantly affect upstream DNA trajectory relative to a cryo-EM structure of RNAPII EC alone (Bernecky et al., 2016). Conversely, Spt5-KOW1 contains an eukaryotic-specific insertion called L1 that increases the extent of upstream DNA contact. These DNA contacts are consistent with increased protection of upstream DNA against exonuclease III digestion by Spt4/5, RfaH, and NusG (Crickard et al., 2016).

**FIGURE 6.**
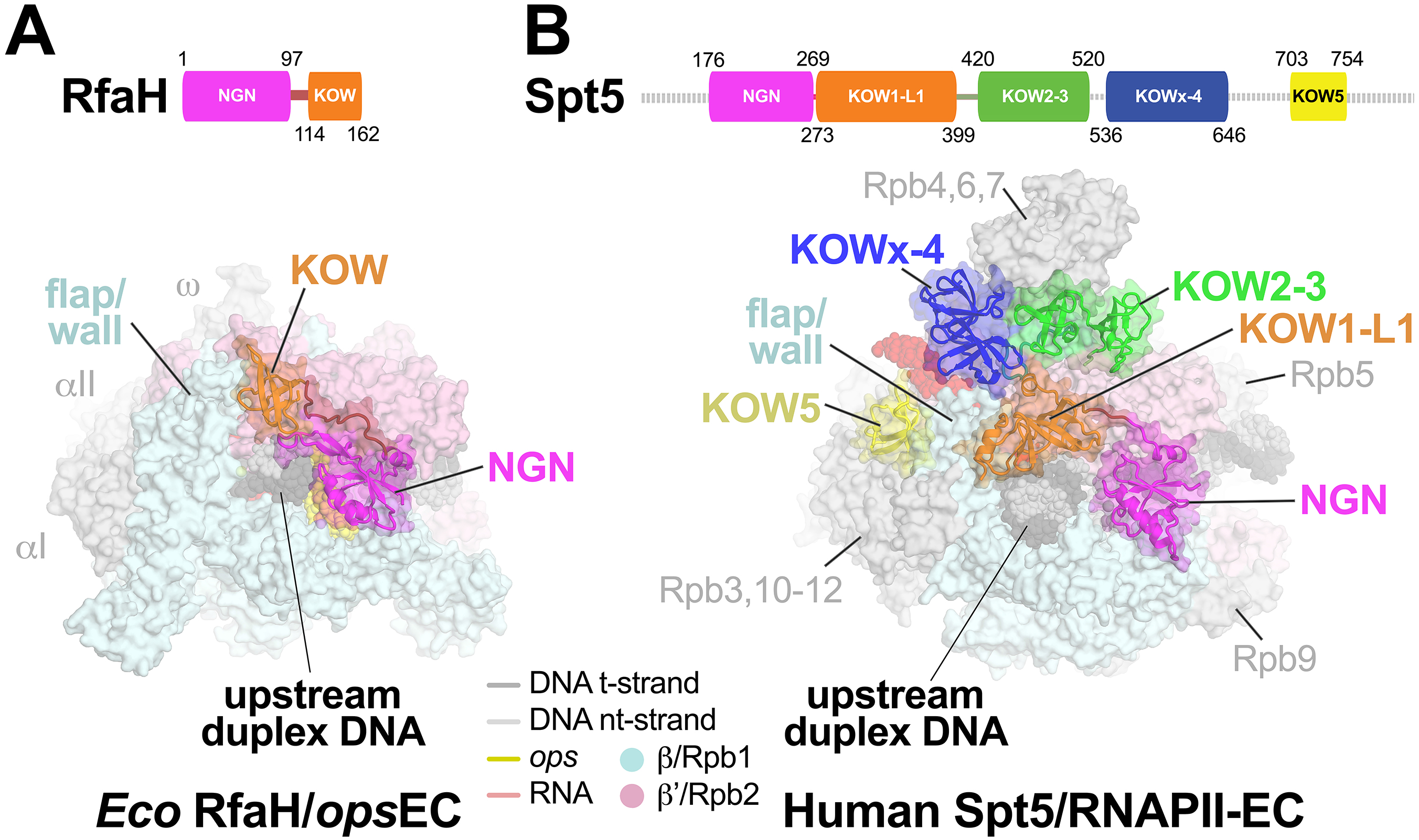
Comparison of the RfaH-*ops*EC with an Spt5-RNAPII-EC. A. (top) Schematic showing RfaH domains (NGN, magneta; KOW, orange), with amino acid residues numbered. (bottom) overall view of the RfaH-*ops*EC structure. The RNAP is shown as a molecular surface, the nucleic acids in CPK format (colored as shown in the key or as labeled). RfaH is shown as a backbone ribbon with a transparent molecular surface, colored as in the schematic above. The RfaH-KOW binds directly above the upstream DNA duplex and next to the β flap/wall. B. (top) Schematic showing the domain organization of Spt5 (Bernecky et al., 2017), with amino acid residues numbered and disordered regions denoted as a dashed line. (bottom) overall view of the human Spt5-RNAPII-EC structure (Bernecky et al., 2017). This structure also includes Spt4, which has been removed for clarity. The RNAP is shown as a molecular surface, the nucleic acids in CPK format (colored as shown in the key or as labeled). Spt5 is shown as a backbone ribbon with a transparent molecular surface, colored as in the schematic above. The Spt5-KOW1-L1 domain binds directly above the upstream DNA duplex and next to the Rbp2 flap/wall. Also see (Ehara et al., 2017).

Both RfaH-KOW and Spt5-KOW1 also affect the RNA exit channel, whose mouth is formed in part by the tip of a module called the flap in bacteria and the wall in eukaryotes. The bacterial flap tip is capped with an α-helix that contacts either *σ* factors or NusA and that is required for their function (Kuznedelov et al., 2002)(Guo et al., 2018). In contrast, the flap tip appears to be dispensable for human RNAPII transcription (Palangat et al., 2011). Both the bacterial flap tip and the eukaryotic wall/flap tip are flexible and often disordered in crystal and cryo-EM structures, but strikingly both are contacted by RfaH-KOW/Spt5-KOW1. These contacts pull the bacterial flap tip away from the RNA exit channel into a novel location (Figure 6). It is possible that this change contributes to RfaH effects on PH-stabilized pausing, since deletion of the flap tip eliminates PH stimulation of pausing (Hein et al., 2014). Given the lack of documented function for the wall, its contcts with Spt5-KOW1 may not have significant effects on transcription. However, the Spt5-KOW2-3 domains bind in the path of the exiting RNA immediately upstream from a bacterial PH, and would partially clash with the location of the PH (Kang et al., 2018). Thus, Spt5 KOW2-3 may directly inhibit formation of nascent RNA structures in the RNAPII RNA exit channel.

## DISCUSSION

Transcription in all cellular organisms is a focal point for the regulation of gene expression. Although the central enzyme of transcription, the multisubunit cellular RNAP, is evolutionarily conserved across all life (Lane and Darst, 2010a; 2010b), this conservation does not extend to regulation. For instance, each RNAP faces similar mechanistic challenges during promoter-specific initiation but relies on evolutionarily unrelated basal factors (Werner and Grohmann, 2011). Similarly, unrelated factors control the elongation and termination phases of the transcription cycle, with one exception. NusG/Spt5 elongation factors are structurally (Figure 6) and functionally homologous, making them the only transcription regulators that are conserved in all domains of life (Werner, 2012). We report here key insights into the mechanistic basis for the regulation of RNAP function by this family of universal regulators from cryo-EM structures of *Eco* NusG and its operon-specific paralog *Eco* RfaH engaged with an EC (Figure 1). We show that (i) NusG and RfaH can suppress backtrack-pausing by stabilizing base-pairing of the upstream duplex DNA (Figure 2); (ii) RfaH achieves operon-specificity in part by specific recognition of an *ops* DNA hairpin in the exposed nt-DNA of the EC transcription bubble (Figure 3), and (iii) RfaH suppresses PH-stabilized pausing by preventing RNAP swiveling (Figures 4, 5), an RNAP conformational change required for the PH effect on prolonging pausing (Kang et al., 2018).

### The mechanistic basis for NusG/RfaH regulation of RNAP pausing

RNAP pausing is a key mechanism for regulating gene expression in all organisms (Mayer et al., 2017; Zhang and Landick, 2016). The mechanistic and structural basis for transcriptional pausing is understood in greatest detail in bacteria (Artsimovitch and Landick, 2000). Pauses initially arise when specific sequences prevent complete translocation of the DNA, resulting in an offline elemental paused EC (ePEC) with posttranslocated RNA transcript but pre-translocated DNA (Kang et al., 2018). This hybrid state resists NTP binding, providing time for the ePEC to isomerize into other, more long-lived paused states: (i) Backtracking prolongs pausing by disengaging the RNA 3’-terminus from the RNAP active site (Nudler, 2012); (ii) PH formation stabilizes the RNAP in the swiveled conformation (Figure 4C) that prolongs pausing by allosterically inhibiting trigger-loop folding require for the optimal active site configuration (Kang et al., 2018). The results presented here allow us to propose mechanistic hypotheses for the complex effects of NusG and RfaH on RNAP elongation.

The primary effects of NusG-NGN and RfaH-NGN on the EC are i) stabilizing base-pairing in the upstream duplex DNA (NusG and RfaH; Figure 2), and ii) inhibiting RNAP swiveling (RfaH; Figures 4C, 5). Formation of the ePEC is associated with an incompletely translocated intermediate without major conformational changes in the RNAP (Kang et al., 2018), and accordingly NusG and RfaH are not known to have strong effects on the ePEC (Larson et al., 2014).

Both NusG and RfaH increase the overall transcription elongation rate by suppressing backtrack pausing (Herbert et al., 2010; Svetlov et al., 2007; Turtola and Belogurov, 2016). A detailed analysis of NusG effects on the individual steps of the RNAP nucleotide addition cycle, on backtracking, and on the conformational stability of the upstream duplex DNA in the EC led (Turtola and Belogurov, 2016) to propose that NusG stabilizes the first upstream bp (the −10 bp in our scaffold; Figure 1A), thereby suppressing backtracking since the −10 bp must melt for backtracking to occur. Our structural and biochemical analyses (Figure 2) support the conclusion that both NusG and RfaH suppress backtracking by stabilizing the −10 bp of the upstream duplex DNA.

Both NusG and RfaH binding are sterically incompatible with RNAP swiveling (Figure 4C), but only RfaH efficiently suppresses PH-stabilized pausing (Figure 5C). The results of the CTR assay (Figures 5E, F) and the NusG/RfaH retention assay (Figures 5G, H) argue that the binding energy of RfaH to the EC is sufficient to inhibit PH formation and suppress RNAP swiveling, whereas NusG binding energy is not. These results explain how RfaH can suppress PH-stabilized pausing by counteracting RNAP swiveling, whereas NusG cannot.

### Adaptations in RfaH confer operon-specificity

In bacteria, the specialized NusG paralog RfaH has maintained key functions of the NGN (EC binding, suppression of backtrack pausing) and tethered KOW (coordinating transcription-translation coupling through ribosome interactions) but has developed an elaborate regulatory mechanism to confer operon-specificity. These include specific recognition of an *ops*-hairpin in the exposed nt-DNA of the transcription bubble (Figure 3A) and a molecular switch that auto-inhibits RfaH-NGN-RNAP interactions in the absence of *ops* recognition (Burmann et al., 2012) (Figure 7B).

**FIGURE 7.**
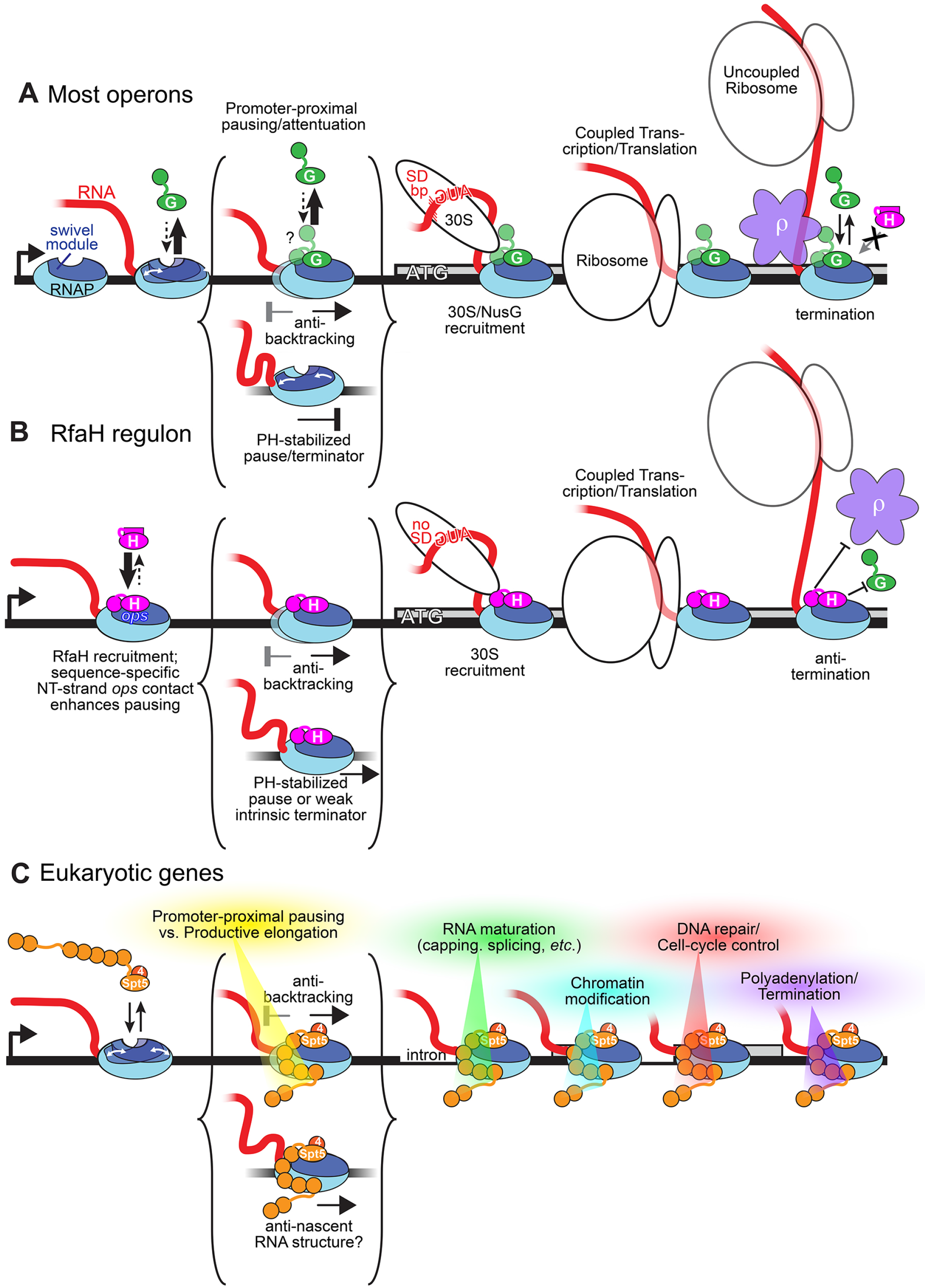
Comparison of NusG, RfaH, and Spt5 action as elongation regulators. A. On most bacterial operons, NusG (G, *green*) associates with RNAP weakly until RNAP enters a protein-coding gene and NusG-KOW interaction with 30S is possible (during initiation or slight uncoupling of transcription and translation). Weak NusG interaction is sufficient to inhibit backtracking but not to inhibit PH-induced RNAP swiveling. When transcription and translation become significantly uncoupled, NusG activates Rho-dependent termination via NusG-KOW-Rho interaction. B. On *ops*-containing operons, RfaH (H, *magenta*) is recruited by contacts to the *ops* hairpin in the exposed nt-DNA (*yellow*; Figure 3A). The refolded RfaH-KOW domain interacts with RNAP and the tightly bound RfaH excludes NusG, inhibits backtracking, and inhibits PH stimulation of pausing by preventing RNAP swiveling (Figure 4C). RfaH associates with 30S similarly to NusG, but inhibits Rho indirectly upon significant transcription-translation uncoupling by excluding NusG. C. Spt4/5 (4 and 5, *orange*) associates with RNAPII in the promoter-proximal region though NusG homologous contacts of its NGN and of 5 of its 7 KOWs (Bernecky et al., 2017; Ehara et al., 2017). Bound Spt5 inhibits backtracking, may inhibit swiveling and formation of nascent RNA structures, and mediates the regulatory switch between promoter-proximal pausing/attenuation and productive elongation in part by affecting actions of NELF and P-TEFb (Kwak and Lis, 2013; Mayer et al., 2017). The full set of Spt5 interactions involved in this switch as well as in downstream roles in mediating RNA maturation, chromatin modifications, recruiting of DNA repair factors, cell-cycle control, polyadenylation, and termination remains incompletely defined (Crickard et al., 2017; Hartzog and Fu, 2013; Meyer et al., 2015).

The cryo-EM structure of the RfaH-*ops*EC represents the RfaH ‘loading’ complex, revealing the structural adaptations that allow specific *ops*-RfaH recognition (Figure 3). The auto-inhibitory RfaH C-terminal helical hairpin sterically blocks the protein-protein interactions between RfaH and the RNAP β’ CH (Figure 4B) but would not interfere with *ops*-RfaH interactions. Presumably, pausing of the EC precisely at *ops* displays the *ops* hairpin in the single-stranded nt-DNA to allow for RfaH recognition and establishment of RfaH/RNAP β subunit interactions (Figure 4B). These partial RfaH-*ops*EC interactions provide sufficient dwell time so that thermal fluctuations of the RfaH auto-inhibitory helical hairpin allow RfaH-RNAP β’ CH interactions, stabilizing the RfaH-NGN-EC interactions and freeing the RfaH-C-terminal domain (CTD) to refold into the KOW (Figure 1B). Unlike the NusG-KOW, which is disordered in our cryo-EM maps, the RfaH-KOW was pinned down through interactions with the EC in a sub-population of the particles (Figures 1B, 6A, S4), similar to the Spt5-KOW1-L1 domain (Figure 6B). Weak RfaH-KOW-EC interactions could help prevent the RfaH-CTD from competing with the RNAP β’ CHs for RfaH-NGN interactions in the absence of other RfaH-KOW interactors, such as the ribosome (Figure 7B).

The RfaH-NGN binds to the EC with a higher affinity than the NusG-NGN (Belogurov et al., 2009) (Figure 5H), a second adaptation that allows RfaH to function in an operon-specfic manner by excluding NusG from the RfaH-EC. Moreover, the tighter binding of the RfaH-NGN confers its ability to counteract RNAP swiveling, providing a mechanistic basis for the ability of RfaH to suppress PH-stabilized pauses (Figures 5C, 7B).

### Complex roles of NusG/Spt5 factors *in vivo*

The *in vivo* roles of both NusG and Spt5 are complex. They both stimulate transcript elongation by RNAP over much of the genome, but more importantly for the cell they serve as recruitment platforms for accessory factors to coordinate transcription elongation with other cellular functions (Figure 7). For example, NusG plays a crucial role in ρ-dependent termination through direct NusG-KOW-ρ interactions (Pasman and Hippel, 2000). In addition, NusG plays critical functional and/or scaffolding roles in multiprotein assemblies that effect rRNA antitermination (Squires et al., 1993; Torres et al., 2004).

RfaH function is confined to transcription units containing an *ops* sequence in an upstream segment but is no less complex. RfaH uses its ability to suppress backtrack and RNA hairpin-stabilized pausing and coordinate with the ribosome (but not ρ) through its KOW to ensure the efficient transcription of long operons.

The control and release of promoter-proximal pausing by RNAPII is a major mechanism for regulating gene expression in metazoans (Kwak and Lis, 2013). Not surprisingly, Spt5 and its heterodimeric partner Spt4 (together called DRB-sensitivity-inducing factor, or DSIF) play important roles in promoter-proximal pausing. DSIF cooperates with NELF to enable promoter-proximal pausing (Wu et al., 2003; Yamaguchi et al., 1999), and is modified by P-TEFb to transition into an elongation activator (Kwak and Lis, 2013). A key question now is if DSIF and its regulatory partners control RNAPII pausing using similar mechanisms as those revealed here for NusG and RfaH.

## AUTHOR CONTRIBUTIONS

Conceptualization, I.A., R.L., S.A.D.; Investigation, J.Y.K., R.A.M., Y.N., J.S., T.V.M.; Analysis, J.Y‥, I.A., R.L., S.A.D.; Writing, J.Y.K., I.A., R.L., S.A.D.; Supervision, I.A., R.L., S.A.D.; Funding Acquisition, I.A., R.L., S.A.D.

## ACKNOWLEDGMENTS

We thank K. Uryu and D. Acehan at The Rockefeller University Electron Microscopy Resource Center for help with EM sample preparation, M. Ebrahim and J. Sotiris at The Rockefeller University Cryo-EM Resource Center and E. Eng at the New York Structural Biology Center for help with data collection, and members of our research groups for helpful comments on the manuscript. Some of the work presented here was conducted at the National Resource for Automated Molecular Microscopy located at the New York Structural Biology Center, supported by grants from the NIH National Institute of General Medical Sciences (GM103310) and the Simons Foundation (349247). This work was supported by National Institute of Health grants R01 GM67153 to I.A., R01 GM38660 to R.L. and R35 GM118130 to S.A.D.

## METHODS

### KEY RESOURCES TABLE

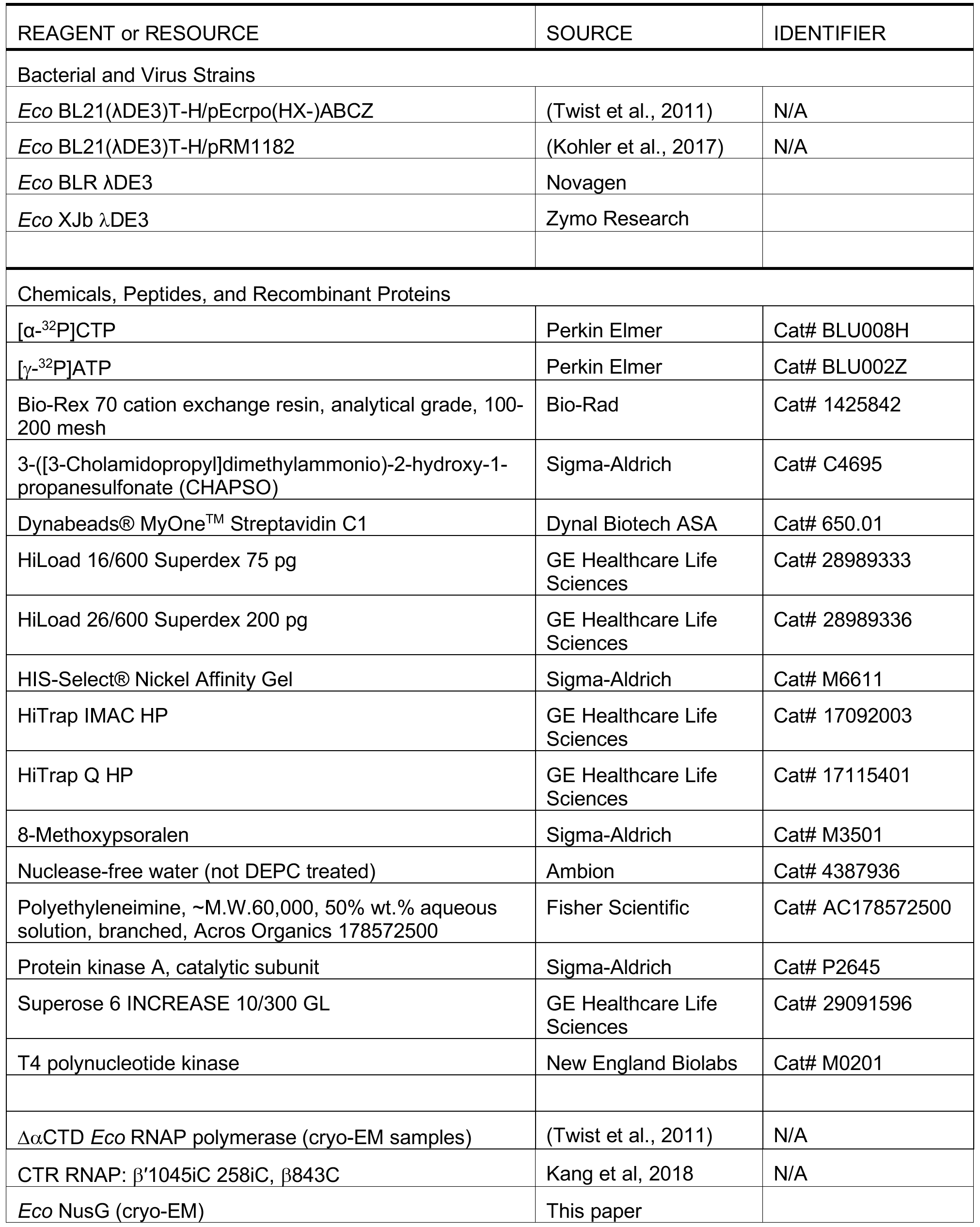

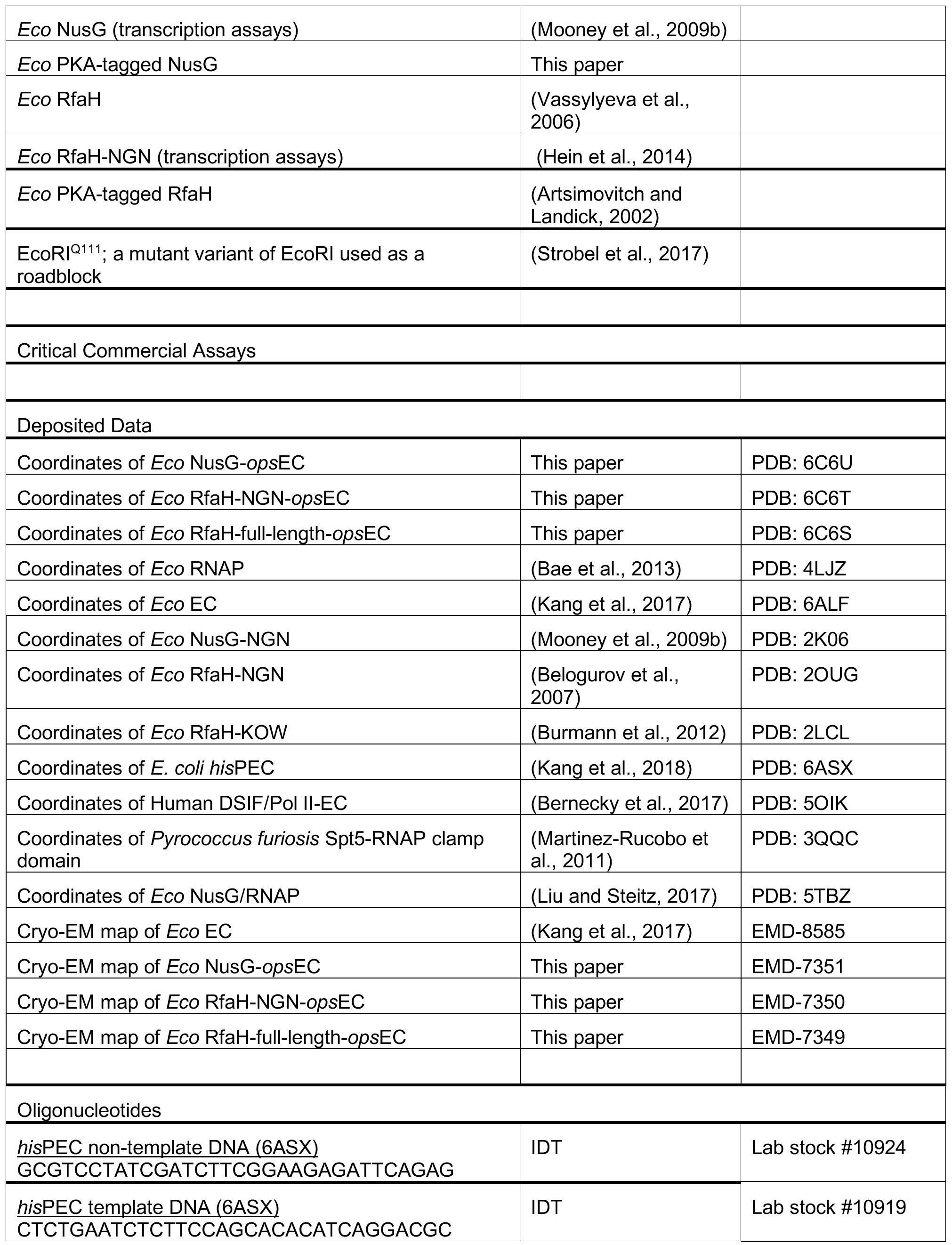

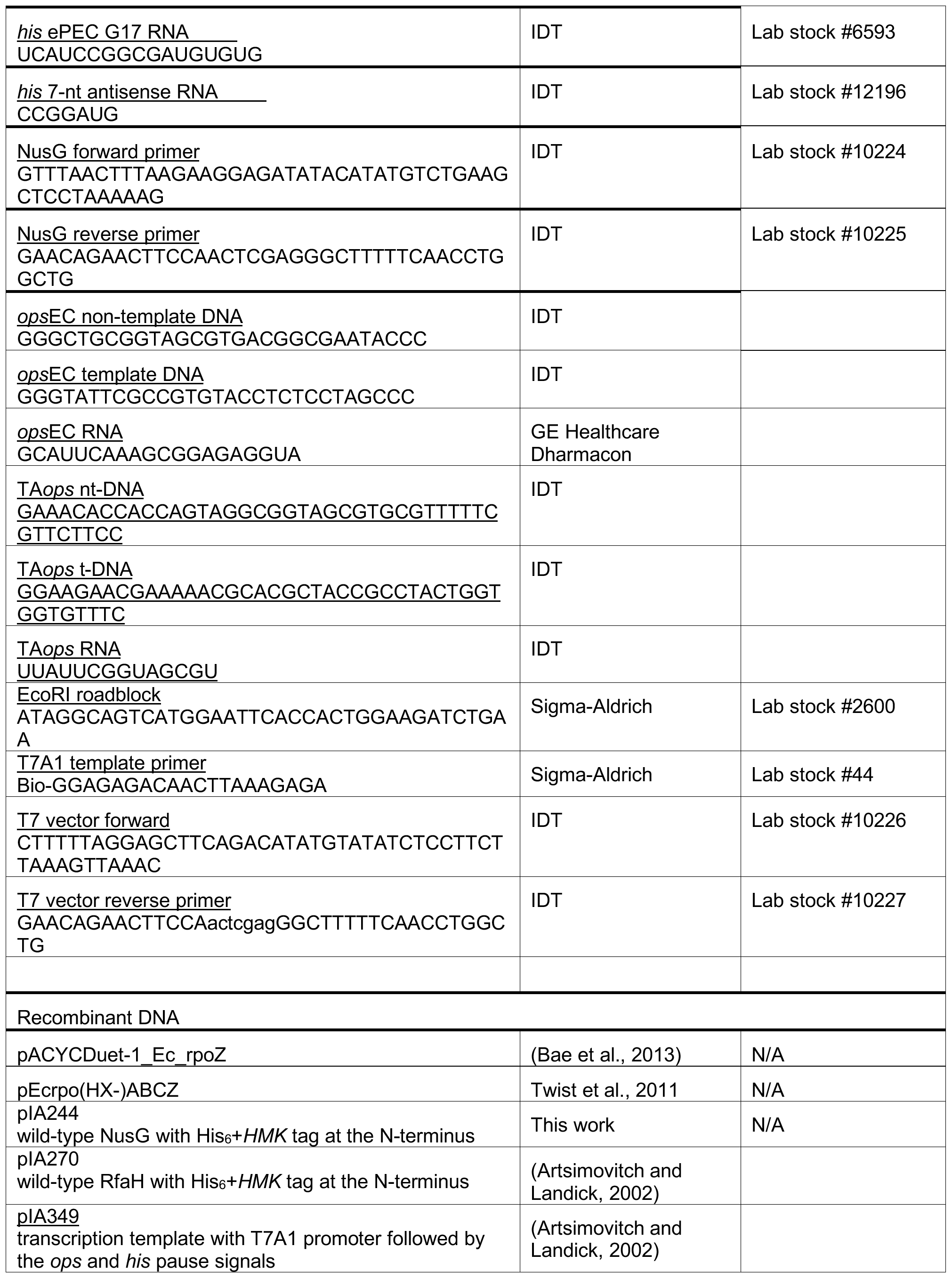

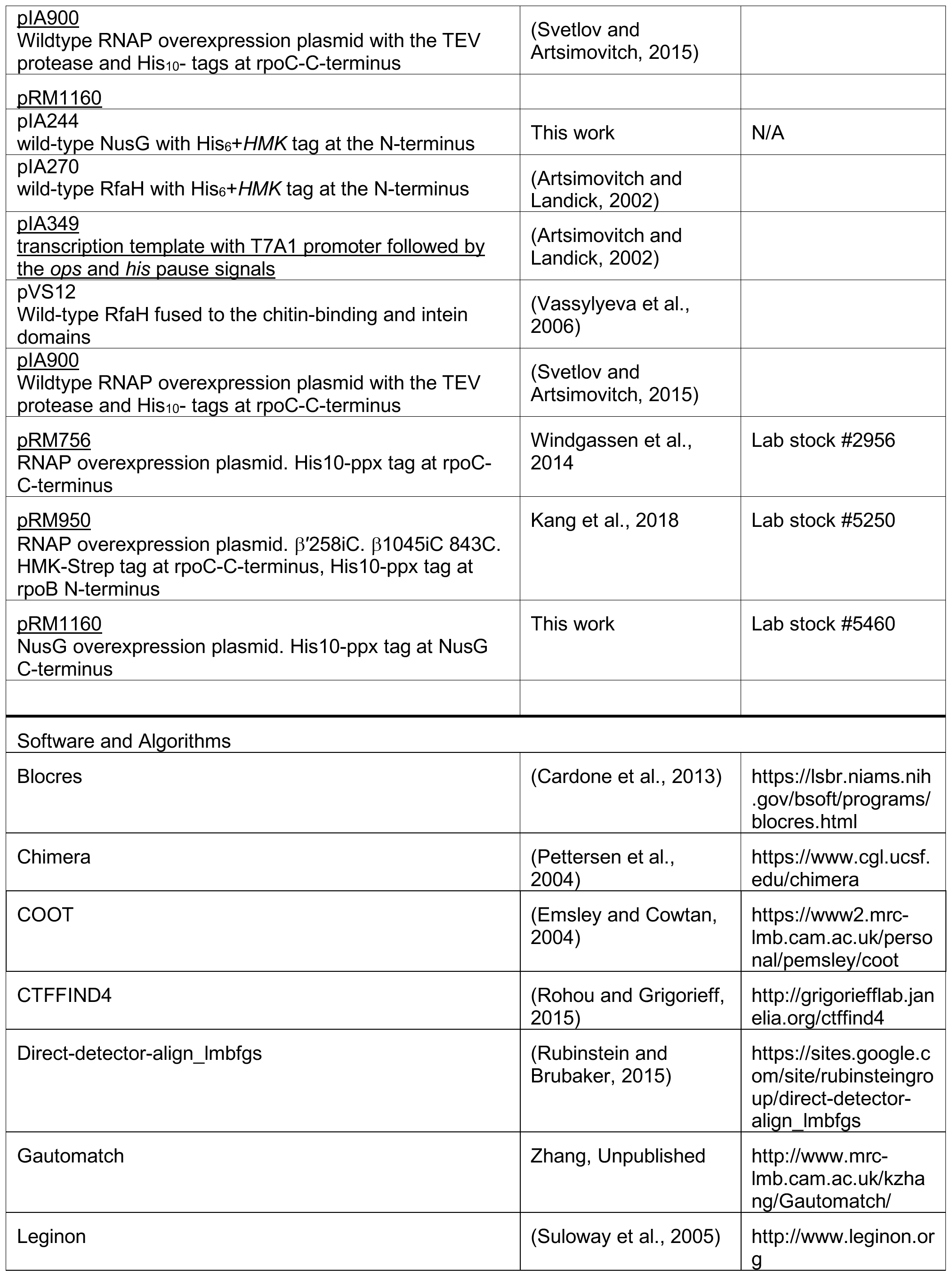

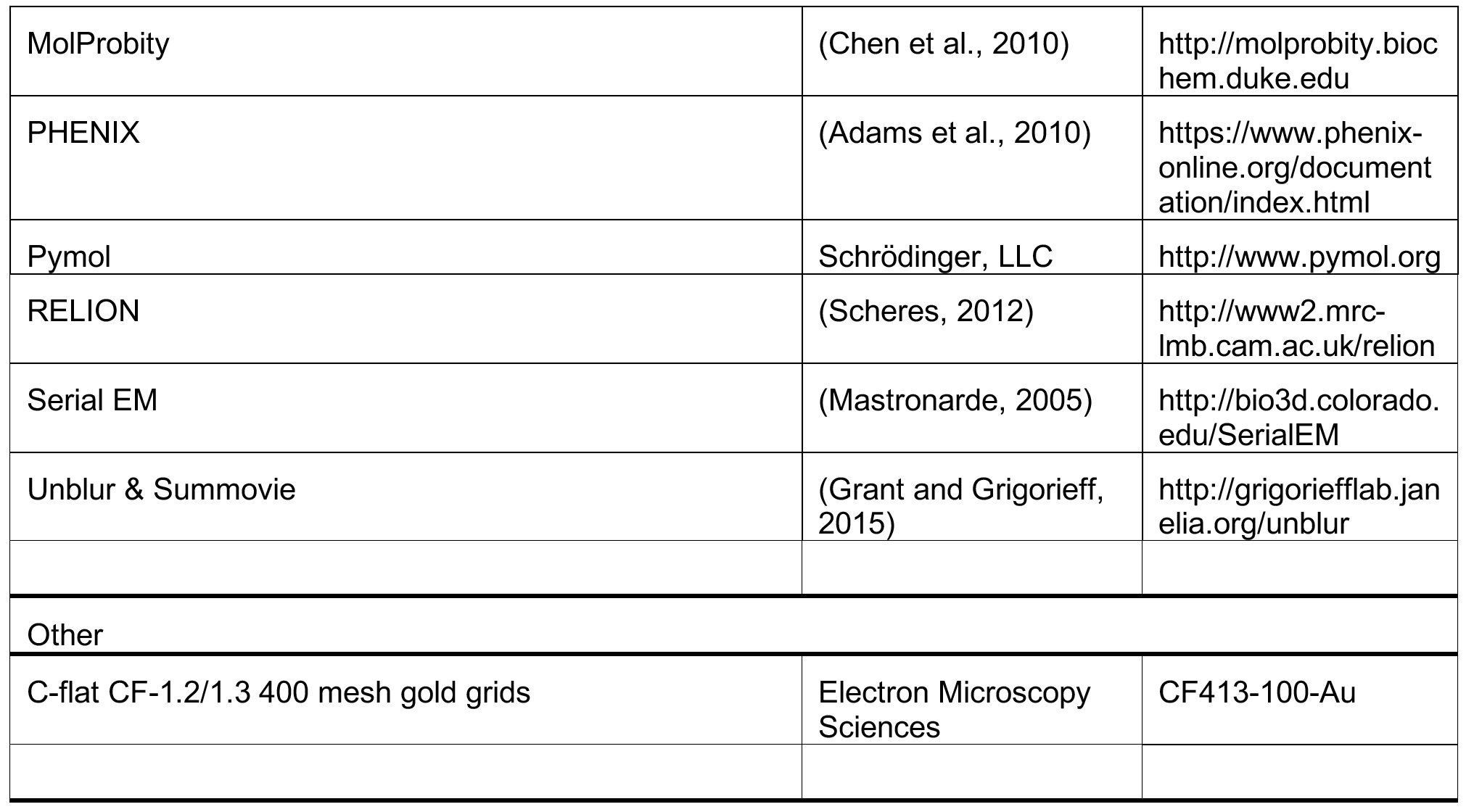

### CONTACT FOR REAGENT AND RESOURCE SHARING

S. A. Darst

## METHODS DETAILS

### RNAP expression and purification for Cryo-EM

*Eco* RNAP lacking the αCTDs was prepared as described previously (Twist et al., 2011). Glycerol was added to the purified RNAP to 15% (v/v), and the sample was aliquoted and flash-frozen in liquid nitrogen. The aliquots were stored at −80 °C until use.

### NusG expression and purification

NusG was purified from pRM1160. pRM1160 was generated by Gibson assembly of the wild-type *nusG* gene PCR amplified from *Eco* chromosomal DNA using primers 10224 and 10225 with a vector fragment amplified from pRM756 using primers 10226 and 10027. pRM1160 is kanamycin-resistant, contains the T7 promoter upstream of *nusG*, and encodes *nusG* followed by a precision protease cleavage site and ten histidine residues.

To prepare full-length *Eco* NusG, plasmids encoding NusG with a C-terminal (His)_6_-tag were grown in *Eco* BL21 (DE3) in LB with 50 μg/ml kanamycin at 37 °C to an OD_600_ of 0.5, induced for protein expression by addition of IPTG to final concentration of 0.5 mM, grown for an additional 3 hours, then harvested by centrifugation. Harvested cells were resuspended in 50 mM Tris-HCl, pH 8.0, 300 mM NaCl, 0.1 mM EDTA, 1 mM β-mercaptoethanol, 0.1 mM PMSF, and lysed in a continuous flow French Press (Avestin, Ottawa, Ontario, Canada). The lysate was centrifuged and the supernatant was loaded onto a HiTrap IMAC column (GE healthcare Life Sciences, Pittsburgh, PA) charged with Ni^2+^. The column was washed with 20 mM Tris-HCl, pH 8.0, 300 mM NaCl, β-mercaptoethanol, 30 mM imidazole, then eluted with an imidazole gradient to 250 mM. The eluted protein was inclubated with human rhinovirus (HRV) 3C protease to cleave the C-terminal (His)_6_-tag and dialyzed for overnight at 4 °C against 20 mM Tris-HCl, pH 8.0, 100 mM NaCl, 1 mM DTT. The dialyzed and tag-cleaved protein was loaded onto a Hitrap Q (GE Healthcare Life Sciences) and eluted with a NaCl gradient. The eluted protein was purified by gel filtration chromatography on a HiLoad Superdex75 column (GE Healthcare Life Sciences) in 10 mM Tris-HCl, pH 8.0, 500 mM NaCl, 5% (v/v) glycerol, 0.5 mM EDTA, 1 mM DTT. Additional glycerol was added to the purified NusG to 15% (v/v), and the sample was aliquoted and flash-frozen in liquid nitrogen. The aliquots were stored at −80 °C until use.

### RfaH expression and purification

Wild-type *Eco* RfaH was expressed and purified as described previously (Vassylyeva et al., 2006), flash-frozen in liquid nitrogen and stored at −80 °C until use.

### Purification and labeling of HMK-tagged RfaH and NusG

Plasmids encoding NusG and RfaH with an N-terminal (His)_6_-tag and a protein kinase A recognition site (RRASV) were grown in *Eco* XJb (DE3) in LB with 40 μg/ml kanamycin at 37 °C to an OD_600_ of 0.4, induced by addition of IPTG to final concentration of 0.2 mM, and grown at 18 °C overnight; L-arabinose was added to 0.07% 2 hours before harvesting. The cells were harvested by centrifugation, resuspended in 50 mM Tris-HCl, pH 6.9, 500 mM NaCl, 5% Glycerol and 1 mM β-mercaptoethanol supplemented with complete, EDTA-free Protease Inhibitor Cocktail (Roche) and incubated with 0.25 mg/ml of Lysozyme on ice for 30-40 minutes. The cells were then lysed by sonication. The lysate was centrifuged and the supernatant was loaded onto a gravity column with Ni Sepharose 6 Fast Flow resin (GE Healthcare Life Sciences, Pittsburgh, PA) equilibrated with 50 mM Tris-HCl, pH 6.9, 500 mM NaCl, 5% Glycerol and 1 mM β-mercaptoethanol. The column was washed 20 mM imidazole. The proteins were eluted with an imidazole gradient to 250 mM in 50 mM Tris-HCl, pH 6.9, 80 mM NaCl, 5% (v/v) Glycerol and 1 mM β-mercaptoethanol, directly loaded onto a 5-ml HiTrap Heparin column, and eluted with an NaCl gradient to 1 M. The peak fractions were dialyzed against 10 mM Tris-HCl, pH 7.9, 250 mM NaCl, 1 mM DTT, 0.1 mM EDTA, 50% (v/v) glycerol and flash-frozen for storage at −80 °C.

### RfaH-*ops*EC and NusG-*ops*EC preparation for Cryo-EM

Synthetic DNA oligonucleotides were obtained from Integrated DNA Technologies (Coralville, IA), RNA oligonucleotides from GE Healthcare Dharmacon (Lafayette, CO). The nucleic acids for the *ops*-scaffold (Figure 1A) were dissolved in RNase-free water (Ambion/ThermoFisher Scientific, Waltham, MA) at 0.2-1 mM. Template DNA and RNA were annealed at a 1:1 ratio in a thermocycler (95 °C for 2 min, 75 °C for 2 min, 45 °C for 5 min, followed by steady cooling to 25 °C at 1 °C/min). The annealed RNA-DNA hybrid was stored at −80 °C until use. Purified *Eco* RNAP was buffer-exchanged over a Superose 6 INCREASE (GE Healthcare Life Sciences) column into 20 mM Tris-HCl, pH 8.0, 120 mM potassium acetate, 5 mM MgCl_2_, 5 mM DTT. The eluted protein was mixed with the pre-annealed RNA-DNA hybrid at a molar ratio of 1:1.3 and incubated for 15 min at room temperature. Nt-DNA and additional 5 mM MgCl_2_ was added and incubated for 10 min. The complex was concentrated by centrifugal filtration (EMD Millipore, Billerica, MA) to 4.0-5.5 mg RNAP/ml concentration before grid preparation.

### Cryo-EM grid preparation

Before freezing, CHAPSO was added to the samples to 8 mM final concentration. C-flat (Protochips, Morrisville, NC) CF-1.2/1.3 400 mesh gold grids were glow-charged for 15 s prior to the application of 3.5 μl of the complex sample (4.0-5.5 mg/ml protein concentration), then plunge-frozen in liquid ethane using a Vitrobot mark IV (FEI, Hillsboro, OR) with 100% chamber humidity at 22 °C.

### Cryo-EM data acquisition and processing for NusG-*ops*EC

The grids were imaged using a 300 keV Titan Krios (FEI) equipped with a K2 Summit direct electron detector (Gatan, Pleasanton, CA). Images were recorded with Leginon (Suloway et al., 2005) in counting mode with a pixel size of 1.07 Å and a defocus range of 0.8 to 2.5 μm (Figure S3A). Data were collected with a dose of 8 electrons/physical pixel/s. Images were recorded with a 10 s exposure and 0.2 s sub-frames (50 total frames) to give a total dose of 69.9 electrons/Å^2^. Structural biology software was accessed through the SBGrid consortium (Morin et al., 2013). Dose fractionated subframes were aligned and summed using Unblur (Grant and Grigorieff, 2015). The contrast transfer function was estimated for each summed image using CTFFIND4 (Rohou and Grigorieff, 2015). From the summed images, particles were automatically picked in Gautomatch (Zhang, unpublished; see Key Resource Table), manually inspected, and then individually aligned using direct-detector-align_lmbfgs software (Rubinstein and Brubaker, 2015). The aligned particles were subjected to 2D classification in RELION specifying 100 classes (Scheres, 2012), and poorly populated classes were removed, resulting in 514,900 particles (Figure S3B). These particles were 3D autorefined in RELION using a map of *Eco* elongation complex (EMD-8585; Kang et al., 2017), low-pass filtered to 60 Å resolution as an initial 3D template. With this initial model, 3D classification was performed without alignment with a soft mask generated in Chimera (Pettersen et al., 2004) and RELION. The soft mask excluded flexible RNAP domains (SI1, SI3, flap tip helix, and single-stranded nucleic acids) of EC. Among the classes from the 3D classification, the best-resolved and most-populated class was 3D autorefined and subjected to the second 3D classification without alignment with the soft mask that was used in the first 3D classification. From this classification, the best-resolved class containing 171,900 particles was 3D autorefined with solvent flattening, and post-processed in RELION, yielding the final reconstruction at 3.7 Å resolution (Figures S2, S3D). Local resolution calculations (Figure S3F) were performed using blocres (Cardone et al., 2013).

### Cryo-EM data acquisition and processing for RfaH:EC

The grids were imaged using a 300 keV Titan Krios (FEI) equipped with a K2 Summit direct electron detector (Gatan, Pleasanton, CA). Images were recorded with Serial EM (Mastronarde, 2005) in super-resolution counting mode with a super resolution pixel size of 0.650 Å and a defocus range of 0.8 to 2.4 μm (Figure S5A). Data were collected with a dose of 8 electrons/physical pixel/s (1.3 Å pixel size at the specimen). Images were recorded with a 15 s exposure and 0.3 s sub-frames (50 total frames) to give a total dose of 71.0 electrons/Å^2^. Dose fractionated subframes were 2 × 2 binned (giving a pixel size of 1.3 Å), aligned and summed using Unblur (Grant and Grigorieff, 2015). The contrast transfer function was estimated for each summed image using CTFFIND4 (Rohou and Grigorieff, 2015). From the summed images, particles were automatically picked in Gautomatch (Zhang, unpublished; see Key Resource Table), manually inspected, and then individually aligned using direct-detector-align_lmbfgs software (Rubinstein and Brubaker, 2015). The aligned particles were subjected to 2D classification in RELION specifying 100 classes (Scheres, 2012), and poorly populated classes were removed, resulting in 389,200 particles (Figure S5B). These particles were 3D autorefined in RELION using a map of *Eco* elongation complex (EMD-8585; Kang et al., 2017), low-pass filtered to 60 Å resolution as an initial 3D template. With this initial model, 3D classification was performed without alignment with a soft mask generated in Chimera and RELION. The soft mask excluded flexible RNAP domains (SI1, SI3, flap tip helix, and single-stranded nucleic acids) of EC. Among the 3D classes, the two best-resolved classes were combined, 3D autorefined, and subjected to second 3D classification without alignment with the soft mask that was used in the first 3D classification. From this classification, the best-resolved class was 3D autorefined with solvent flattening, and post-processed in RELION, yielding the final reconstruction at 3.5 Å resolution (Figures S4, S5D).

To resolve heterogeneity around RfaH the upstream duplex DNA, focused 3D classification on the unmodeled region was performed (Scheres, 2012). A soft map containing RNAP, nucleic acids, and the RfaH-NGN was generated in Chimera and RELION to make a subtracted particle stack in RELION. The subtracted particles were 3D classified into six classes without alignment, reverted to the original (unmasked) particles, and 3D autorefined. Among the classes, one class resolved the RfaH-KOW domain and flap-tip helix. The particles in this class were post-processed, yielding the final reconstruction at 3.7 Å resolution (Figure S4). Local resolution calculations (Figure S5G) were performed using blocres (Cardone et al., 2013).

### Model building, refinement and validation

To build initial models, *Eco* core enzyme (PDB ID 4LJZ with σ^70^ removed; (Bae et al., 2013) and nucleic acids (PDB ID 6ALF; Kang et al., 2017) were fitted into the electron density maps using Chimera. NusG-NGN (PDB ID 2K06; (Mooney et al., 2009b), RfaH-NGN (PDB ID 2OUG; (Belogurov et al., 2007), and RfaH-KOW (PDB ID 2LCL; (Burmann et al., 2012) were also fitted into NusG-*ops*EC, RfaH-NGN-*ops*EC, and focused RfaH-*ops*EC cryo-EM maps accordingly. These initial models were real-space refined against the working half map using Phenix real-space-refine (Adams et al., 2010). In the refinement, domains in the core and nucleic acids were rigid-body refined, then subsequently refined with secondary structure restraints. At the end of refinement, Fourier shell correlations (FSC) were calculated between the refined model and the half map used for refinement (work), the other half map (free), and the full map to assess over-fitting (Figures S3E, S5E, S5F).

### Psoralen crosslinking

Scaffolds were assembled from synthetic TA*ops* oligonucleotides, t-DNA was end-labeled with [γ^32^P]-ATP using T4 polynucleotide kinase (PNK; NEB). Following labeling, oligonucleotides were purified using QIAquick Nucleotide Removal Kit (Qiagen). To assemble a scaffold, RNA and t-DNA oligonucleotides were combined in PNK buffer and annealed in a PCR machine as follows: 5 min at 45 °C; 2 min each at 42, 39, 36, 33, 30, and 27 °C, 10 min at 25 °C. 12 pmoles of t-DNA/RNA hybrid were mixed with 14 pmoles of His-tagged core RNAP in 30 μl of TB [20 mM Tris-HCl, pH 7.9, 5% (v/v) Glycerol, 40 mM KCl, 5 mM MgCl_2_, 10 mM β-mercaptoethanol], and incubated at 37 °C for 10 min. 15 βl of His-Select^®^ HF Nickel Affinity Gel (Sigma-Aldrich) was washed once in TB and incubated with 20 μg Bovine Serum Albumin in a 40-μl volume for 15 min at 37 °C, followed by a single wash step in TB. The t-DNA/RNA/RNAP complex was mixed with the Affinity Gel for 15 min at 37 °C on a thermomixer (Eppendorf) at 900 rpm, and washed twice with TB. 30 pmoles of the nt-DNA oligonucleotide were added, followed by incubation for 20 min at 37 °C, one 5-min incubation with TB supplemented with 1 M KCl in a thermomixer, and five washes with TB. The assembled ECs were eluted from beads with 90 mM imidazole in a 15-μl volume, purified through a Durapore (PVDF) 0.45 μm Centrifugal Filter Unit (Merck Millipore), and resuspended in TB. For crosslinking, the ECs were supplemented with 6.3% (v/v) DMSO and 0.92 mM 8- methoxypsoralen and incubated for 2 min at 37 °C, followed by addition of 50 nM RfaH, 500 nM NusG, or storage buffer, and a 3-min incubation at 37 °C. Complexes were then exposed to 365 nm UV light (8W Model UVLMS-38; UVP, LLC) for 20 min on ice. The reactions were quenched with an equal volume of Stop buffer (8 M Urea, 20 mM EDTA, 1 × TBE, 0.5 % Brilliant Blue R, 0.5 % Xylene Cyanol FF). Samples were heated for 2 min at 95 °C and separated by electrophoresis in denaturing acrylamide (19:1) gels (7 M Urea, 0.5X TBE). The gels were dried and the products were visualized and quantified using a FLA9000 Phosphorimaging System (GE Healthcare), ImageQuant Software, and Microsoft Excel.

### Cys Triplet Reporter (CTR) assays

Nucleic-acid scaffolds used to reconstitute *his*PEC for Cys triplet reporter cross-linking assays (Figure 5E) were assembled on purified DNA and RNA oligonucleotides as described (Hein et al., 2014). Briefly, 10 μM RNA, 12 μM template DNA, and 15 μM nt-DNA (Key Resource Table) were annealed in reconstitution buffer (RB; 20 mM Tris-HCl, pH 7.9, 20 mM NaCl, and 0.1 mM EDTA). To assemble complexes, scaffold (2 μM) was mixed with limiting CTR RNAP (1 μM; CTR RNAP: β′1045iC 258iC, β843C) in transcription buffer (50 mM Tris-HCl, pH 7.9, 20 mM NaCl, 10 mM MgCl_2_, 0.1 mM EDTA, 5% glycerol, and 2.5 μg of acetylated bovine serum albumin/ml). NusG-NGN or RfaH-NGN proteins were added to 1 μM and combined with cystamine and DTT to final concentrations of 2.5 mM and 2.8 mM, respectively, to generate a redox potential of −0.36. Reactions were incubated for 60 min at room temperature and then stopped by addition of iodoacetamide to 15 mM. The formation of cysteine cross-links was then evaluated by non-reducing SDS-PAGE (4-15% gradient Phastgel; GE Life Sciences) as described previously (Nayak et al., 2013). Gels were stained with Coomassie Blue and imaged with a CCD camera. The fraction cross-linked was quantified with ImageJ software (Schneider et al., 2012). The experimental error was determined as the standard deviation of measurements from three or more independent replicates.

### RNAP pause assays

The nucleic-acid scaffold (Figure 5A; Key Resources Table) used to reconstitute ECs for pause assays (Figures 5C, D) was assembled as previously described (Hein et al., 2014). Briefly, PAGE-purified G17 RNA (2 nt upstream of the pause site, 10 μM), t-DNA (15 μM), and nt-DNA (20 μM) were annealed in reconstitution buffer (RB; 10 mM Tris-HCl, pH 7.9, 40 mM KCl, and 5 mM MgCl_2_). Scaffolds (2 μM) were incubated with 0.5 μM RNAP for 15 min at 37 °C in Elongation Buffer (EB; 25 mM HEPES-KOH, pH 8.0, 130 mM KCl, 5 mM MgCl_2_, 1 mM dithiothreitol, DTT, 0.15 mM EDTA, 5% glycerol, and 25 μg of acetylated bovine serum albumin/ml) to form ECs, diluted to 0.1 μM in EB, then incubated with heparin (0.1 mg/ml final) for 3 min at 37 °C, and labeled by incorporation of 2 μM [α-^32^P]CMP for 1 min at 37 °C. NusG-NGN or RfaH-NGN (or EB for the ±asRNA conditions) were added to the C18 complexes and incubated at 37 °C for 10 min before adding 7-mer asRNA or equal volume TE for the minus asRNA condition, and then incubated for another 10 min at 37°C to form an RNA duplex mimic of the *his*PEC hairpin. ECs were then assayed for pause-escape kinetics by addition of 100 μM UTP and 10 μM GTP in EB at 37 °C. Reaction samples were removed at time points and quenched with an equal volume of 2X urea stop buffer (8 M urea, 50 mM EDTA, 90 mM Tris-borate buffer, pH 8.3, 0.02% each bromophenol blue and xylene cyanol). All active PECs were then chased out of the pause by addition of 1 mM GTP for 1 min at 37 °C to aid quantitation. RNAs in each quenched reaction sample were separated on a 15% urea-PAGE gel. The gel was exposed to a PhosphorImager screen, and the screen was scanned using Typhoon PhosphorImager software and quantified in ImageQuant (GE Life Sciences). The fraction of RNA at pause (U19) as a function of time was fit to single-or double-exponential decay functions using KaleidaGraph to obtain the amplitudes of bypass, slow pause, and slower pause species and pause escape rates.

### Retention of RfaH and NusG on the EC

A linear template was generated by PCR of pIA349 using a top biotinylated primer and a bottom primer with an EcoRI recognition site. The template (8 pmoles) was incubated with EcoRI^Q111^ (3 μM; to achieve complete occupancy) in 40 μl BB (20 mM Tris-HCl, pH 7.9, 6% glycerol, 50 mM KCl, 5 mM MgCl_2_, 1 mM β-mercaptoethanol) for 15 min at 37°C. To form an immobilized halted G37 EC, holo-RNAP (8 pmoles), ApU (100 μM) and 5 μM each CTP, GTP and ATP were added together with 20 μl of prewashed Streptavidin coated magnetic beads (Dynabeads^®^ MyOne^TM^ Streptavidin C1) and incubated for 15 min at 37 °C. The halted complexes were washed three times with 500 μl of BB using a Magnetic Separation Stand (Promega). UTP was added at 5 μM for 5 min at 37 °C, followed by three washes. Then ATP, GTP, and CTP were added at 5 μM to form G42 (*ops*10) EC. The sample was divided into two aliquots; to one, ^32^P-labeled RfaH was added to 50 nM, and to the other ^32^P-labeled NusG was added to 470 nM, followed by a 5-min incubation at 37°C and three washes. Each reaction was split again into three aliquots: a) no further treatment; b) 5-min chase at 37°C with 100 μM NTPs; and c) 5-min chase at 37°C with 100 μM NTPs and 5 μM unlabeled NusG. After three washes with BB, samples were measured in a LS6500 Multi-Purpose Scintillation Counter (Beckman Coulter). The experiment was done in triplicates.

### Accession numbers

The cryoEM density maps have been deposited in the EM Data Bank with accession codes EMD-7351 (NusG-*ops*EC), EMD-7350 (RfaH-NGN-*ops*EC), and EMD-7349 (RfaH-full-length-*ops*EC). Atomic coordinates have been deposited in the Protein Data Bank with accession codes 6C6U (NusG-*ops*EC), 6C6T (RfaH-NGN-*ops*EC), and 6C6S (RfaH-full-length-*ops*EC).

## SUPPLEMENTAL INFORMATION

Supplemental Information includes 6 figures and 1 table and can be found with this article online at‥.

**Table S1.**
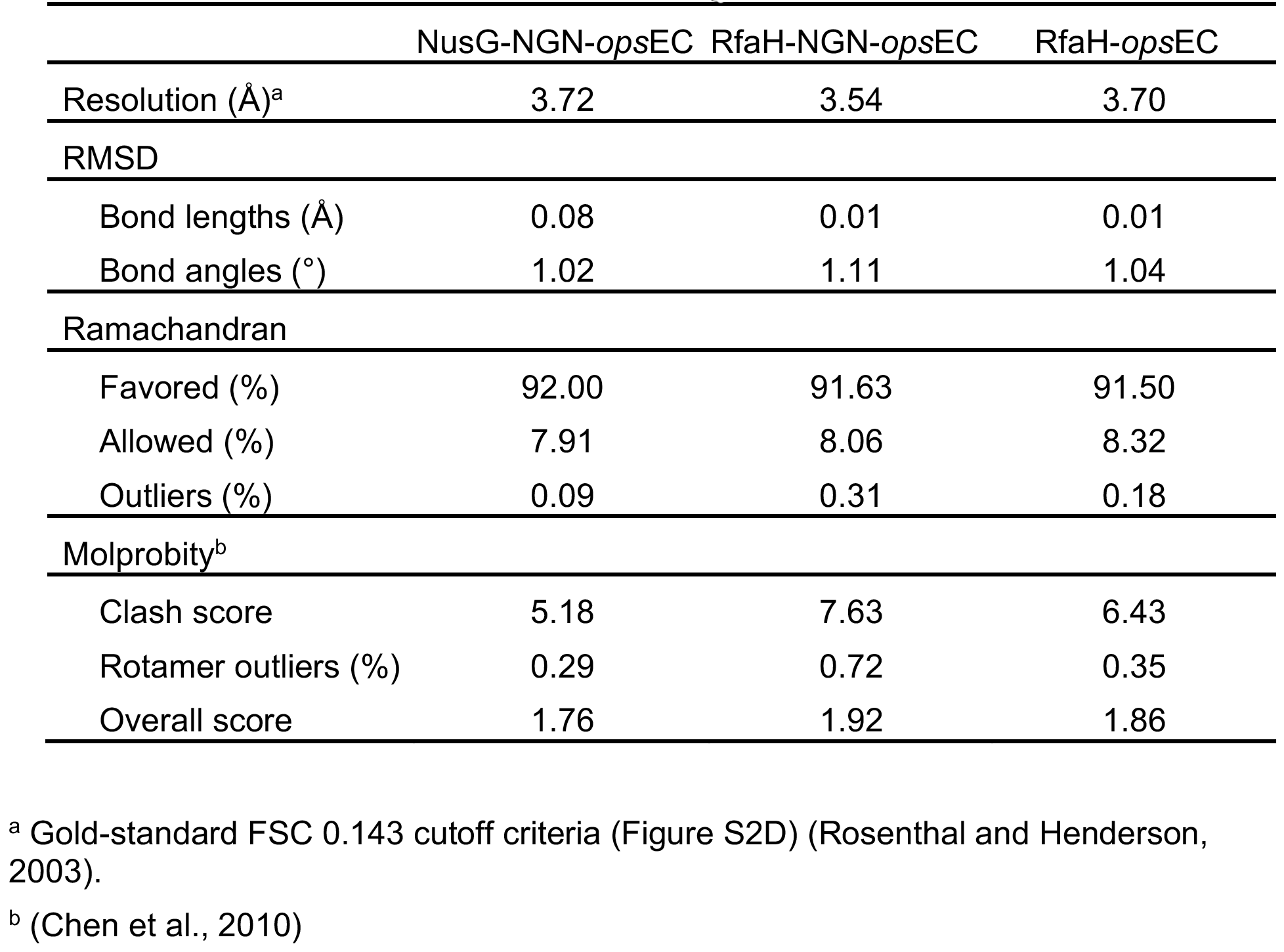
Refinement statistics. Related to Figure 1.

**FIGURE S1.**
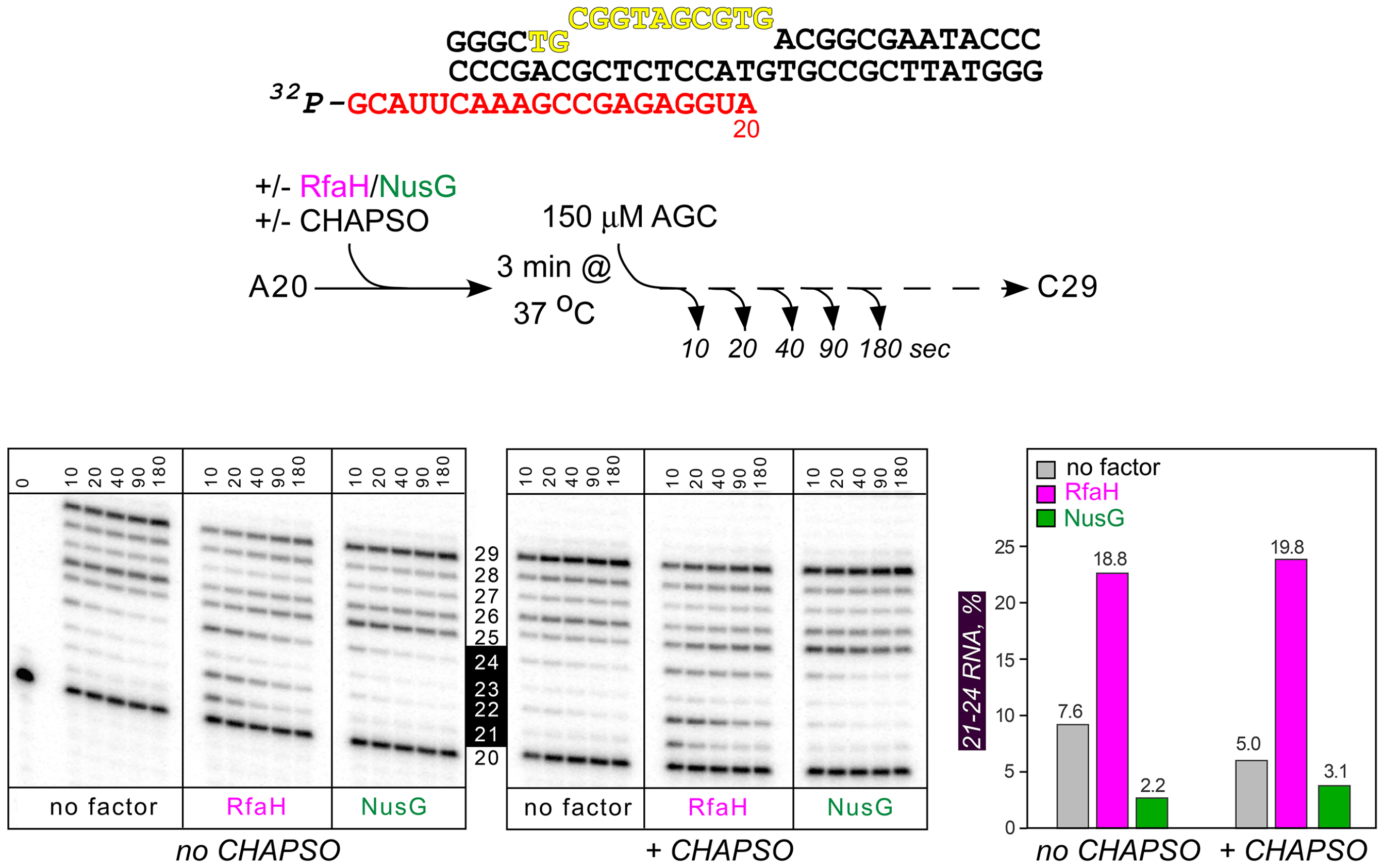
*Ops* scaffold verification. Related to Figure 1. NusG and RfaH effects on RNA chain elongation on the scaffold used for cryo-EM experiments (shown on top). Scaffolds were assembled with the ^32^P-labeled RNA strand in the cryo-EM buffer and preincubated with RfaH (100 nM) or NusG (1000 nM), in the absence or in the presence of 8 mM CHAPSO. Elongation was restarted upon addition of 0.15 mM ATP, GTP, and CTP, aliquots were withdrawn at the indicated times and analyzed on 12% denaturing urea-acrylamide gels (19:1) in 0.5× TBE. Positions of RNA products are indicated, with the region encompassing C21, A22, C23 and G24 RNAs highlighted in black. RfaH and NusG display their characteristic effects immediately downstream from the ops site (A20). Consistent with the patterns observed on standard transcription templates, RfaH promotes RNAP pausing in the 21-24 region, whereas NusG decreases pausing in this region. The bar graph shows the fraction of C21-G24 RNAs (as % of total RNA) at the 40-sec time point. The addition of CHAPSO has only a minor effect on RNAP elongation and response to NusG and RfaH.

**Figure S2.**
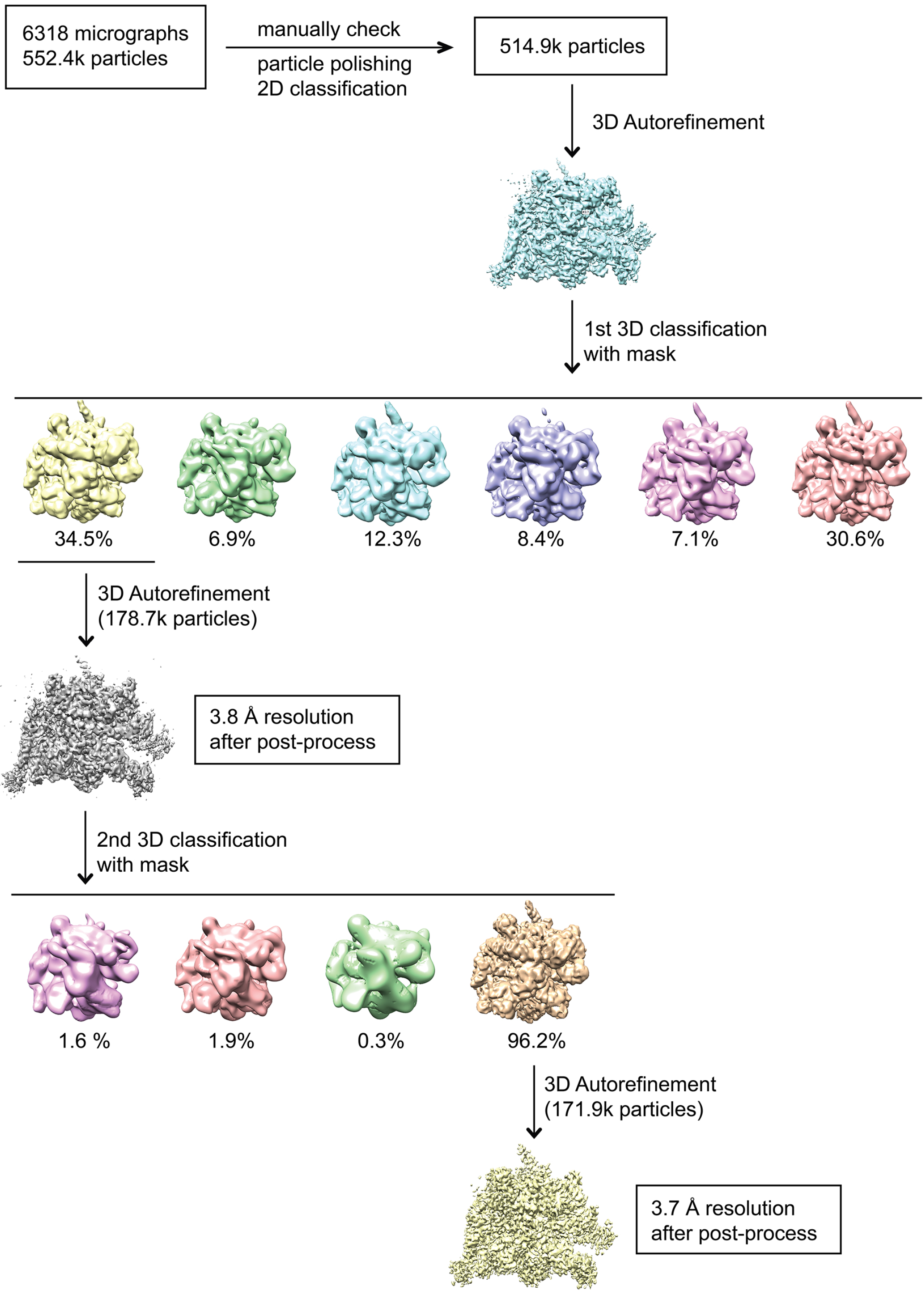
Data processing pipeline for the NusG-*ops*EC cryo-EM data. Related to Figure 1. Flowchart showing the image processing pipeline for the cryo-EM NusG-*ops*EC data starting with 6,318 dose-fractionated movies collected on a 300 keV Titan Krios (FEI) equipped with a K2 Summit direct electron detector (Gatan). Movie frames were aligned and summed using Unblur (Grant and Grigorieff, 2015). Particles were autopicked with Gautomatch (http://www.mrc-lmb.cam.ac.uk/kzhang/Gautomatch/) and manually revised from these summed images. The revised particles were polished using direct-detector-align_lmbfgs (Rubinstein and Brubaker, 2015) for subsequent 2D classification using RELION (Scheres, 2012). After 2D classification, the dataset contained 514,900 aligned particles. These particles were auto-refined in RELION using a model of the *Eco* EC (PDB ID 6ALF; (Kang et al., 2017) as an initial 3D template. 3D classification into six classes was performed on the particles using the refined model and alignment angles. The best class (containing the most particles and having the highest resolution) was subjected to a second 3D classification into four classes. After the second 3D classification, one class containing 33.4% of the starting particles (171,900 particles) was autorefined and post-processed in RELION, yielding the final reconstruction at 3.7 Å resolution.

**Figure S3.**
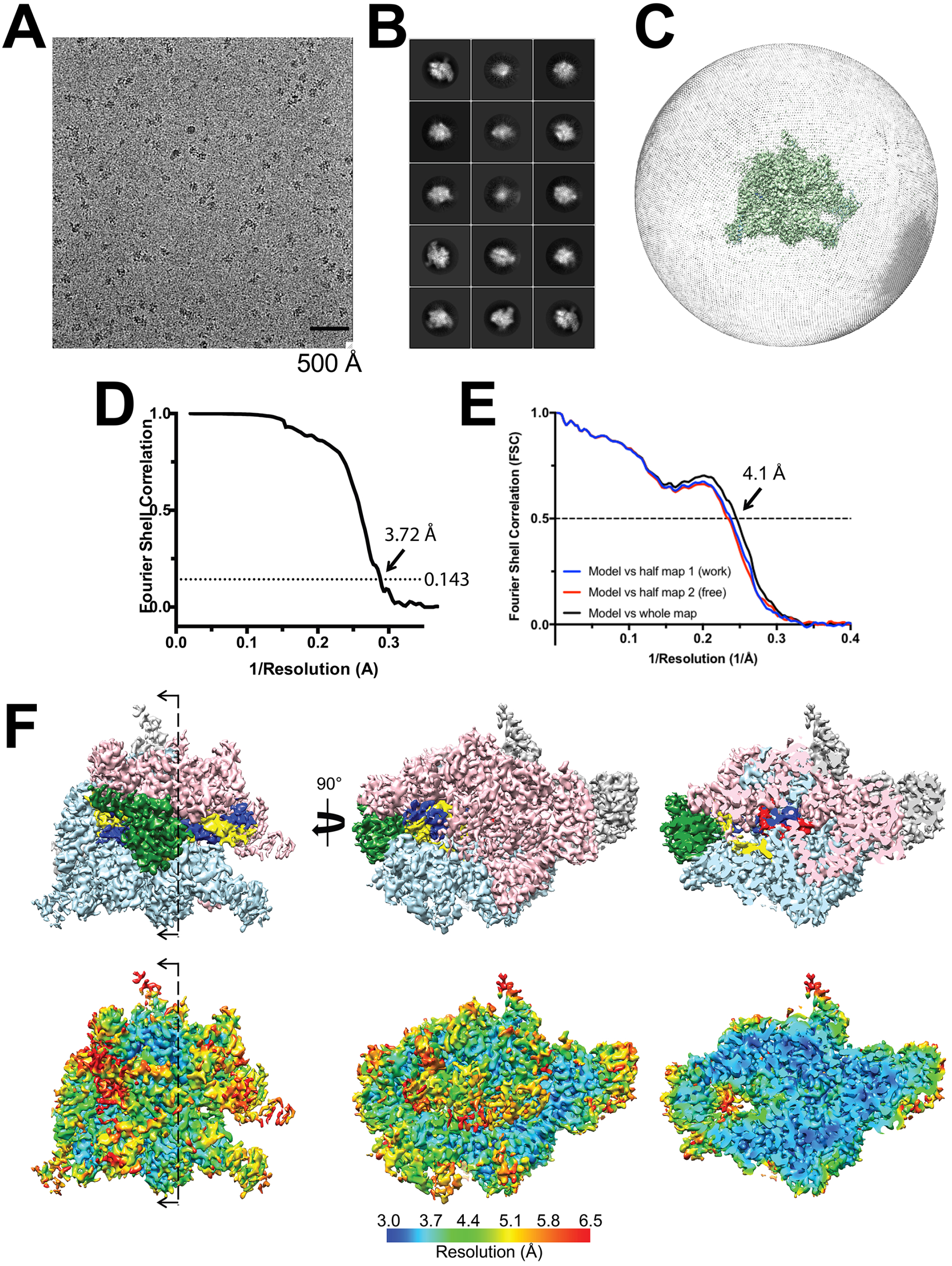
Cryo-EM of the NusG-*ops*EC. Related to Figure 1. A. Representative micrograph of the NusG-*ops*EC in vitreous ice. B. The fifteen most populated classes from 2D classification. C. Angular distribution for NusG-*ops*EC particle projections. D. Gold-standard FSC of the NusG-*ops*EC. The gold-standard FSC was calculated by comparing the two independently determined half-maps from RELION. The dotted line represents the 0.143 FSC cutoff, which indicates a nominal resolution of 3.7 Å. E. FSC calculated between the refined structure and the half map used for refinement (work), the other half map (free), and the full map. F. (top) The 3.7-Å resolution cryo-EM density map of the NusG-*ops*EC is colored as follows: αI, αII, ω subunits, gray; β, cyan; β’, pink; NusG, green; t-DNA, dark blue; nt-DNA, yellow; RNA, red. The rightmost view is sliced as indicated in the leftmost view. (bottom) Same views as (top) but colored by local resolution (Cardone et al., 2013).

**Figure S4.**
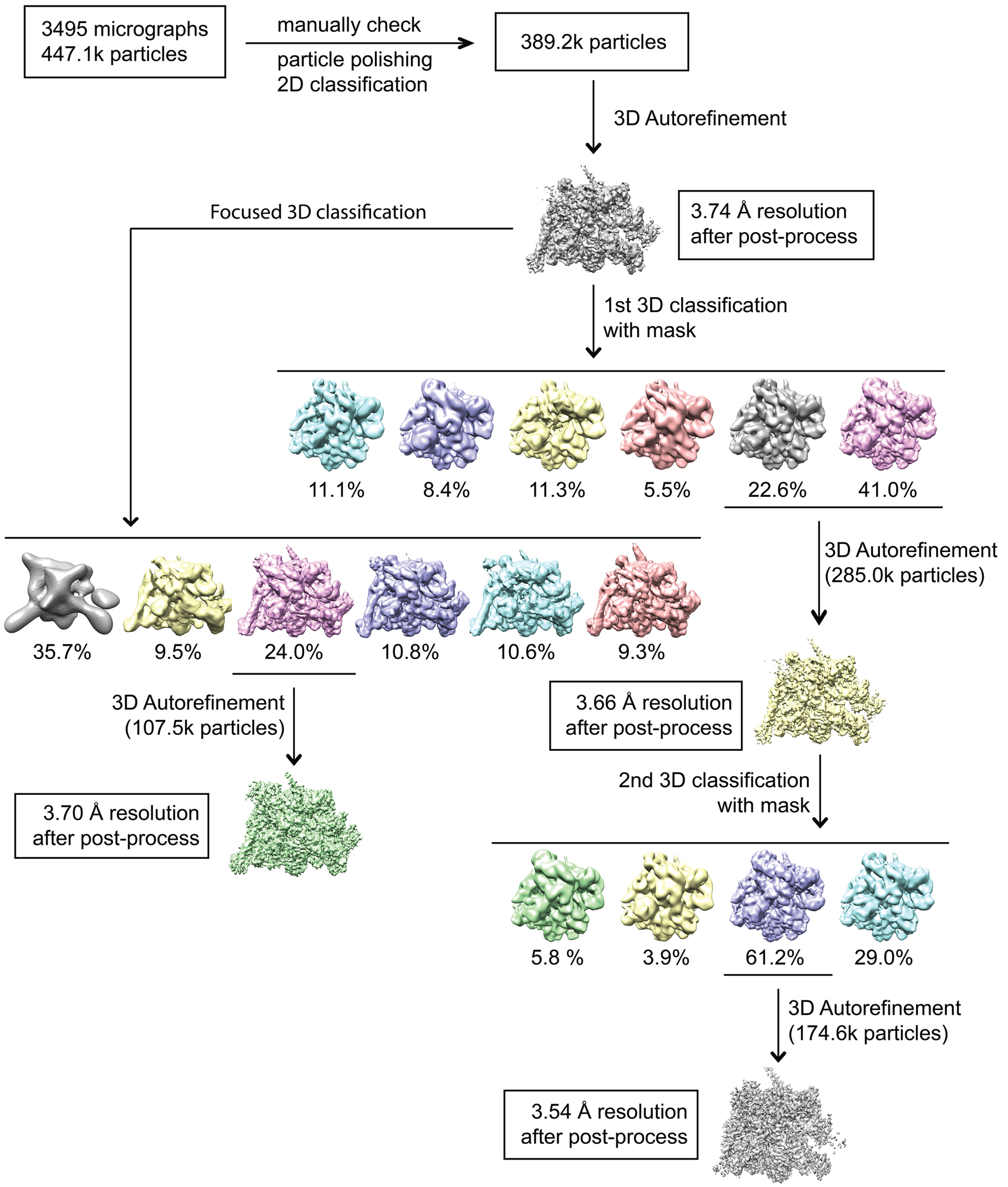
Data processing pipeline for the RfaH-*ops*EC cryo-EM data. Related to Figure 1. Flowchart showing the image processing pipeline for the cryo-EM RfaH-*ops*EC data starting with 3,495 dose-fractionated movies collected on a 300 keV Titan Krios (FEI) equipped with a K2 Summit direct electron detector (Gatan). Movie frames were aligned and summed using Unblur (Grant and Grigorieff, 2015). Particles were autopicked with Gautomatch (http://www.mrc-lmb.cam.ac.uk/kzhang/Gautomatch/) and manually revised from these summed images. The revised particles were polished using direct-detector-align_lmbfgs (Rubinstein and Brubaker, 2015) for subsequent 2D classification using RELION (Scheres, 2012). After 2D classification, the dataset contained 389,200 aligned particles. These particles were auto-refined in RELION using a model of the *Eco* EC (PDB ID 6ALF; (Kang et al., 2017) as an initial 3D template. 3D classification into six classes (right branch) was performed on the particles using the refined model and alignment angles. The best two classes (containing the most particles and having the highest resolutions) was subjected to a second 3D classification into four classes. After the second 3D classification, one class containing 44.9% of the starting particles (174,600 particles) was autorefined and post-processed in RELION, yielding the final reconstruction at 3.5 Å resolution. For the focused 3D classification (left branch of the flowchart), a soft mask that excluded the upstream duplex DNA and nearby protein regions was generated using Chimera and RELION. The mask was used to make a subtracted particle stack in RELION with the filtered map generated in the initial autorefinement. The subtracted particles were 3D classified into six classes without alignment. Among the six classes, one class containing 24% of the starting particles (107,500 particles) had density for the RfaH-KOW domain. The original (unmasked) particles in this class were autorefined and post-processed in RELION, yielding the final reconstruction at 3.7 Å resolution.

**Figure S5.**
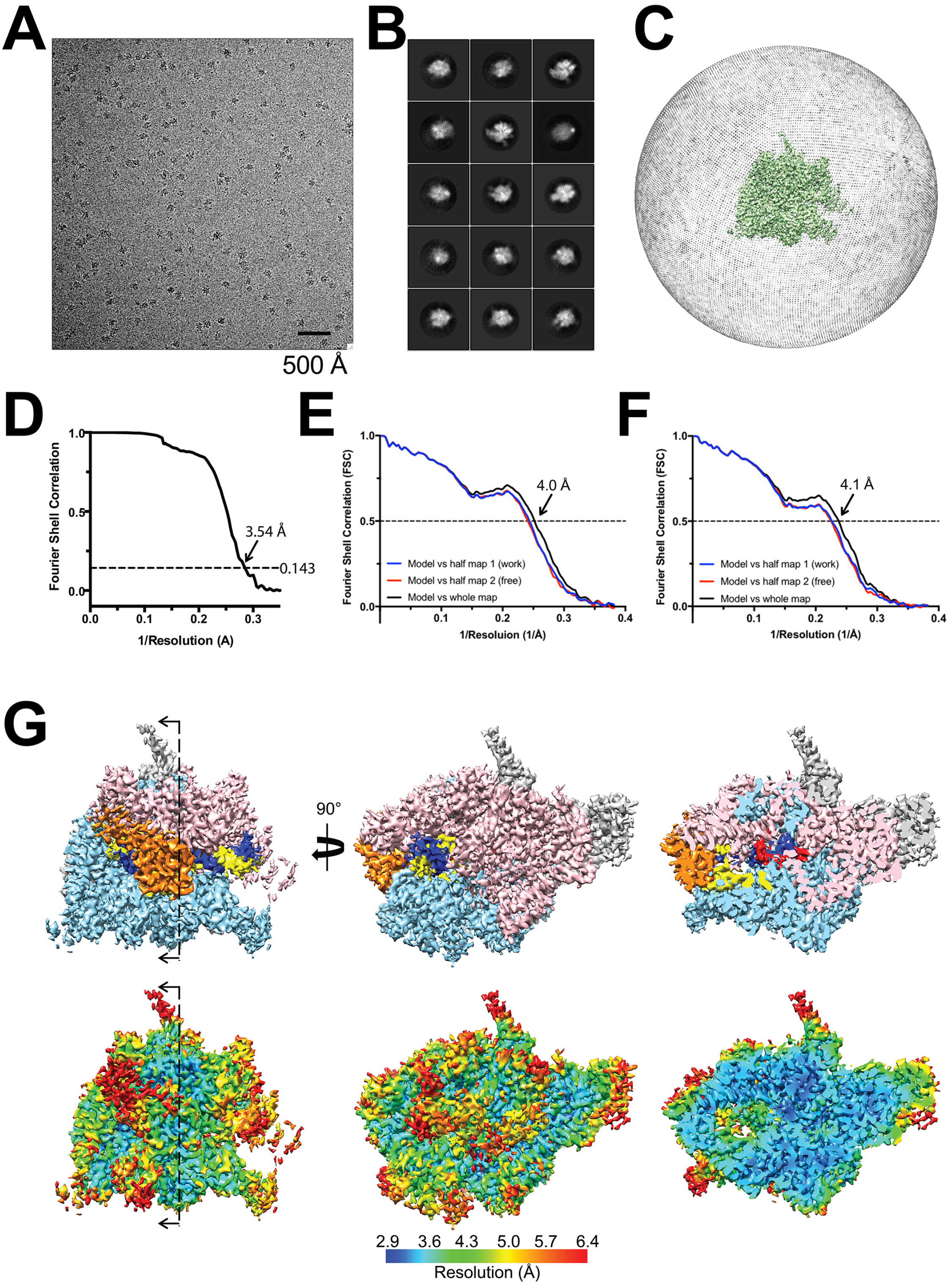
Cryo-EM of the RfaH-*ops*EC. Related to Figure 1. A. Representative micrograph of the RfaH-*ops*E in vitreous ice. B. The fifteen most populated classes from 2D classification. C. Angular distribution for RfaH-*ops*EC particle projections. D. Gold-standard FSC of the RfaH-*ops*EC. The gold-standard FSC was calculated by comparing the two independently determined half-maps from RELION. The dotted line represents the 0.143 FSC cutoff, which indicates a nominal resolution of 3.5 Å. E. RfaH-NGN-*ops*EC: FSC calculated between the refined structure and the half map used for refinement (work), the other half map (free), and the full map. F. RfaH-full-length-*ops*EC: FSC calculated between the refined structure and the half map used for refinement (work), the other half map (free), and the full map. G. (top) The 3.5-Å resolution cryo-EM density map of the RfaH-*ops*EC is colored as follows: αI, αII, ω subunits, gray; β, cyan; β’, pink; RfaH, orange; t-DNA, dark blue; nt-DNA, yellow; RNA, red. The rightmost view is sliced as indicated in the leftmost view. (bottom) Same views as (top) but colored by local resolution (Cardone et al., 2013).

**Figure S6.**
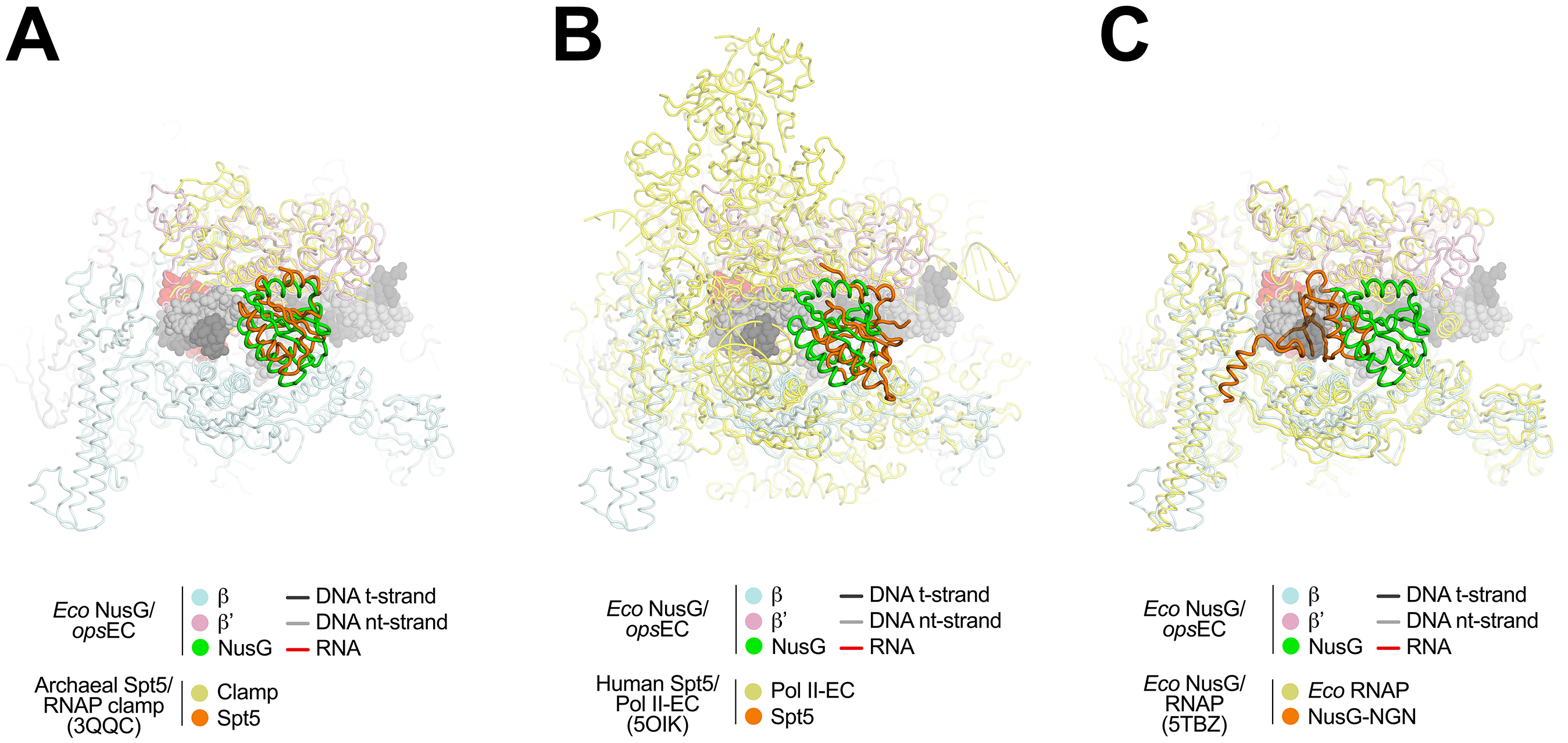
Structural comparisons. Related to Figure 6. Previously determined structures of NusG/Spt5 factors bound to the archaeal RNAP clamp domain (A; 3QQC; (Martinez-Rucobo et al., 2011), bound in a RNAPII-EC (B; 5OIK; (Bernecky et al., 2017), or bound to *Eco* core RNAP (C; 5TBZ; (Liu and Steitz, 2017)) are superimposed with the NusG-*ops*EC via RNAP α-carbons to compare the resulting dispositions of the NusG/Spt5. The color-coding in each panel is indicated in the color keys below. A. The archaeal (*Pyrococcus furiousus*) RNAP clamp domain was superimposed with the *Eco* RNAP from the NusG-*ops*EC, resulting in an rmsd for the 40 α-carbons of the CHs of 1.23 Å. The positions of the NusG-NGN (green) and Spt5 (orange) are shifted relative to each other by about 2.8 Å but the orientation of the domain is conserved. B. The Rpb1 and Rpb2 subunits of human RNAPII from the human DSIF(Spt4/Spt5)/RNAPII-EC were superimposed with the β’ and β subunits (respectively) from the NusG-*ops*EC, resulting in an rmsd for the 40 α-carbons of the CHs of 1.52 Å. The positions of the NusG-NGN (green) and Spt5 (orange) are shifted relative to each other by about 7.9 Å but the orientation of the domain is conserved. C. The *Eco* core RNAP from the NusG/RNAP was superimposed with the RNAP from the NusG-*ops*EC, resulting in an rmsd for the 40 α-carbons of the CHs of 1.94 Å. The orientation of overall fold of the NusG-NGN (orange) is not consistent with the NusG-NGN from the NusG-*ops*EC (green).

